# Breeding with Major and Minor Genes: Genomic Selection for Quantitative Disease Resistance

**DOI:** 10.1101/2021.05.20.444894

**Authors:** Lance F. Merrick, Adrienne B. Burke, Xianming Chen, Arron H. Carter

## Abstract

Most disease resistance in plants is quantitative, with both major and minor genes controlling resistance. This research aimed to optimize genomic selection (GS) models for use in breeding programs needing to select both major and minor genes for resistance. In this experiment, stripe rust (*Puccinia striiformis* Westend. f. sp. *tritici* Erikss.) of wheat (*Triticum aestivum* L.) was used as a model for quantitative disease resistance. The quantitative nature of stripe rust is usually phenotyped with two disease traits, infection type and disease severity. We compared two types of training populations composed of 2,630 breeding lines phenotyped in single plot trials from four years (2016-2020) and 475 diversity panel lines from four years (2013-2016), both across two locations. We also compared the accuracy of models with four different major gene markers and genome-wide association (GWAS) markers as fixed effects. The prediction models used 31,975 markers replicated 50 times using 5-fold cross-validation. We then compared the GS models with marker-assisted selection to compare the prediction accuracy of the markers alone and in combination. The GS models had higher accuracies than marker-assisted selection and reached an accuracy of 0.72 for disease severity. The major gene and GWAS markers had only a small to zero increase in prediction accuracy over the base GS model, with the highest accuracy increase of 0.03 for major markers and 0.06 for GWAS markers. There was a statistical increase in accuracy by using the disease severity trait, the breeding lines, population type, and by combing years. There was also a statistical increase in accuracy using major markers within the validation sets as the mean accuracy decreased. The inclusion of fixed effects in low prediction scenarios increased accuracy up to 0.06 for GS models using significant GWAS markers. Our results indicate that GS can accurately predict quantitative disease resistance in the presence of major and minor genes.

## INTRODUCTION

Plant breeding programs select and improve both qualitative and quantitative traits. Qualitative traits are controlled by a few large-effect genes that are readily detectable and follow a Mendelian inheritance (Chen, 2013). In contrast, quantitative traits are controlled by many small-effect genes that are difficult to distinguish and controlled by quantitative trait loci (QTL; Bernardo, 2008). The genetic control of a trait determines the types of selection that will be most effective for improvement. However, disease resistance can be either a qualitative or a quantitative trait, and therefore varies on the most effective method of improvement (Poland and Rutkoski, 2016). Breeding for disease resistance is a major goal for most breeding programs because of the effect disease has on yield and quality performance.

Breeding for qualitative disease resistance is controlled by one or two large-effect alleles, called resistance (R) genes and further referred to as major genes (Agrios, 2005). Qualitative disease resistance generally follows a race-specific resistance and quickly degrades due to the rapid evolution of new pathogen races (Chen, 2005). Major gene pyramiding can reduce the possibility of major genes’ breakdown by combining multiple major genes for more durable resistance to multiple pathogen races into a single line. Pyramiding is implemented through marker-assisted selection (MAS) and has been an effective method for various crops (Wang et al., 2001; Pietrusińska et al., 2011; Bai et al., 2012; Jiang et al., 2012; Liu et al., 2016b; Singh et al., 2017; Wang et al., 2017). The successful implementation of major genes relies on identifying useful sources of the genes, finding linked markers, confirming the effect in different genetic backgrounds, and finally deploying said major genes (Bernardo, 2008). Implementation is further complicated when it comes to selecting multiple major genes simultaneously for gene pyramiding. A large population is needed to screen and select lines with more than one gene in early generations while still maintaining enough lines to select for other traits in later generations (Poland and Rutkoski, 2016). The difficulty can be further attributed to unfavorable linkage and multiple major gene sources (Bernardo, 2008).

Breeding for quantitative resistance conferred by minor-effect genes tends to produce more durable resistance in breeding lines because it relies on multiple small-effect alleles. Breeding for quantitative resistance requires multiple breeding cycles to improve resistance gradually (Poland and Rutkoski, 2016). The breeding method for quantitative resistance is similar to the methodology used for other complex traits such as grain yield (Rutkoski et al., 2014; Poland and Rutkoski, 2016; González-Camacho et al., 2018). Selecting for quantitative resistance can be completed throughout the breeding process, but disease resistance is commonly completed in earlier generations in order to select for other traits further in the program, similar to qualitative resistance. Therefore, selecting for quantitative resistance in early generations can be difficult due to the lack of replication and environments. However, selecting for resistance in later generations reduces genetic gain due to selection for other traits (Poland and Rutkoski, 2016). Both methods, therefore, reduce the effectiveness of breeding quantitative resistance. One such trait that displays both qualitative and quantitative resistance is stripe rust, also called yellow rust (Yr), caused by *Puccinia striiformis* Westend. f. sp. *tritici* Erikss.).

Stripe rust is one of the most devastating diseases of wheat (*Triticum aestivum* L.) and is highly destructive in the western United States (Chen, 2005; González-Camacho et al., 2018; Liu et al., 2019). Stripe rust can cause more than 90% yield losses in fields planted with susceptible cultivars (Liu et al., 2020). The use of resistant varieties and fungicide applications are the primary methods to control stripe rust (Chen and Line, 1995b; Liu et al., 2020). Stripe rust resistance is categorized into qualitative all-stage resistance (ASR) and quantitative adult-plant resistance (APR).

All-stage resistance is conferred by race-specific genes that are inherited qualitatively with a life span of around 3.5 years per gene (Case et al., 2014; Chen and Kang, 2017). There are over 300 identified QTL conferring resistance to stripe rust (Wang and Chen, 2017). The large number of major genes identified shows numerous resistance alleles available for breeding purposes in various varieties and populations. Previously major genes *Yr5* and *Yr15* have been shown to be effective against all races of the stripe rust pathogen in the United States (Wang and Chen, 2017). However, virulence to *Yr5* has been demonstrated in a few countries not including the United States (Wellings et al., 2009; Zhang et al. 2020; Kharouf et al., 2021; Tekin et al. 2021). Virulence to *Yr15* has only been documented in Afghanistan (Gerechter-Amitai et al., 1989). The virulence to these genes demonstrates the need to not rely on any single major gene to provide resistance in a cultivar.

Adult-plant resistance is usually a non-race-specific quantitative resistance that is associated with durable resistance with some genes being effective for more than 60 years (Chen, 2013). APR is often affected by temperature and also can be referred to as high-temperature adult-plant (HTAP) resistance, which is often controlled by more than one gene mainly with additive effect (Chen and Line, 1995b, 1995a; Liu et al., 2019). HTAP resistance is influenced by the temperature and age of the plants. As the temperature increases, the plant becomes more resistant, and rust development slows down (Chen, 2005). However, in order to confirm HTAP, greenhouse studies with different temperature ranges need to be conducted (Chen, 2005). HTAP and APR resistance are conferred by different loci with varying effects and often display partial resistance, making them difficult to incorporate into new cultivars (Chen and Line, 1995b; Liu et al., 2019). Consequently, APR or HTAP resistance must be improved over multiple selection cycles as mentioned previously (Rutkoski et al., 2014; Poland and Rutkoski, 2016; González-Camacho et al., 2018). APR is generally expressed in later stages of wheat, whereas ASR is expressed throughout the lifecycle of the plant (Wang and Chen, 2017). Therefore it is difficult to identify APR genes due to the masking of their effect by ASR genes. The masking of ASR genes and the quantitative nature of APR result in much of the APR resistance in a population being uncharacterized. It is recommended to combine both ASR and APR genes to take advantage of both types of resistance limitations (Wang and Chen, 2017). The lack of ASR durability coupled with the challenge in identifying and breeding APR creates a unique opportunity for genomic selection. In addition, major genes for ASR are known to interact with APR and accounting for these markers as fixed effects have increased prediction accuracy (Bernardo, 2014; Rutkoski et al., 2014; Arruda et al., 2016).

In many crops, the difficulty in selecting for qualitative and quantitative disease resistance (similar to stripe rust) creates an opportunity for genomic selection (GS) to integrate quantitative resistance by accounting for small effect alleles in the presence of large-effect major genes without the development and analysis of mapping populations and techniques (Poland and Rutkoski, 2016). The goal of this study was to identify the best genomic selection method to select for disease resistance in the presence of both major and minor genes. Stripe rust of wheat was used as an example, as most plant breeders try to capture both ASR and APR’s additive effects simultaneously. Identified GS approaches will be a valuable tool for breeders to facilitate cultivar and parental selection for accumulating favorable alleles for disease resistance in the presence of major and minor resistance genes (Rutkoski et al., 2014; Michel et al., 2017).

## MATERIAL AND METHODS

### Phenotypic Data

Two training populations were used to compare the inclusion of fixed effect markers in populations with different frequencies of stripe rust genes. The first training population consists of F3:5 and doubled-haploid soft white winter wheat breeding lines (BL) developed by the Washington State University (WSU) winter wheat breeding program. The BL population was evaluated for stripe rust in unreplicated single plot trials in Pullman and Lind, Washington planted in 2016, 2017, 2018, and 2020 growing seasons (Table 1). 2019 was not included due to the lack of adequate disease severity in our trials. The BL population was previously selected for stripe rust resistance in headrow plots the year previous to unreplicated trials. Susceptible breeding lines in headrow plots were culled and not included in the BL population which represents a prior selected, closely related breeding line population with similar pedigree sources of stripe rust resistance. The second training population consists of diverse association mapping panel (DP) trials evaluated in unreplicated trials in Central Ferry and Pullman, Washington from 2013 to 2016 (Table 1). The mapping panel consists of varieties and breeding lines from at least six soft white winter wheat breeding programs in the Pacific Northwestand represents diverse backgrounds with potential sources of stripe rust resistance.

**Table 1.**
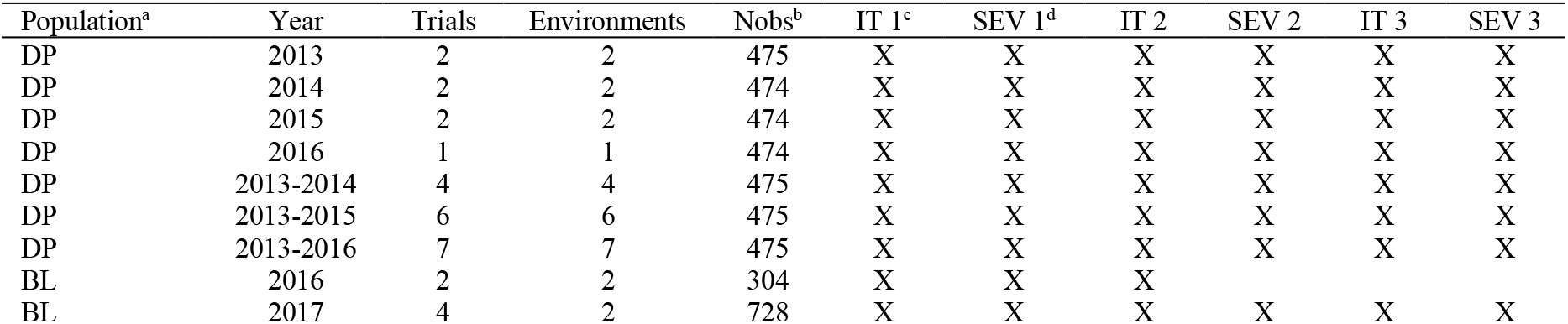
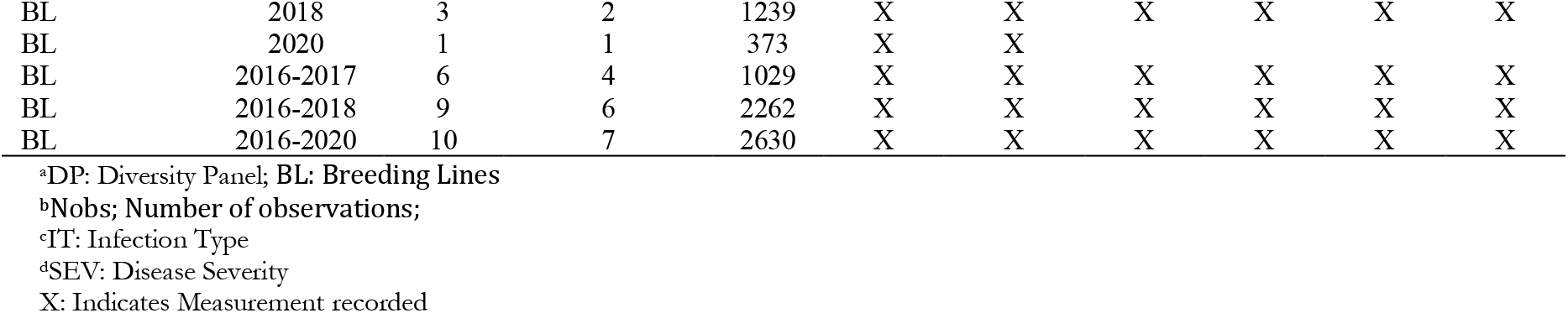
Training populations for stripe rust infection type and disease severity in Central Ferry, Lind, and Pullman, WA from 2013 to 2020.

| Population<sup>a</sup> | Year | Trials | Environments | Nobs<sup>b</sup> | IT 1<sup>c</sup> | SEV 1<sup>d</sup> | IT 2 | SEV 2 | IT 3 | SEV 3 |
| --- | --- | --- | --- | --- | --- | --- | --- | --- | --- | --- |
| DP | 2013 | 2 | 2 | 475 | X | X | X | X | X | X |
| DP | 2014 | 2 | 2 | 474 | X | X | X | X | X | X |
| DP | 2015 | 2 | 2 | 474 | X | X | X | X | X | X |
| DP | 2016 | 1 | 1 | 474 | X | X | X | X | X | X |
| DP | 2013-2014 | 4 | 4 | 475 | X | X | X | X | X | X |
| DP | 2013-2015 | 6 | 6 | 475 | X | X | X | X | X | X |
| DP | 2013-2016 | 7 | 7 | 475 | X | X | X | X | X | X |
| BL | 2016 | 2 | 2 | 304 | X | X | X |  |  |  |
| BL | 2017 | 4 | 2 | 728 | X | X | X | X | X | X |
| BL | 2018 | 3 | 2 | 1239 | X | X | X | X | X | X |
| BL | 2020 | 1 | 1 | 373 | X | X |  |  |  |  |
| BL | 2016-2017 | 6 | 4 | 1029 | X | X | X | X | X | X |
| BL | 2016-2018 | 9 | 6 | 2262 | X | X | X | X | X | X |
| BL | 2016-2020 | 10 | 7 | 2630 | X | X | X | X | X | X |

<sup>a</sup>DP: Diversity Panel; BL: Breeding Lines
<sup>b</sup>Nobs; Number of observations;
<sup>c</sup>IT: Infection Type
<sup>d</sup>SEV: Disease Severity
X: Indicates Measurement recorded

The disease traits measured were stripe rust infection type (IT) and severity (SEV). The recordings of these traits were dependent on natural infection and stripe rust incidence at the time of observation and were not previously inoculated. Some trials had three observations for stripe rustand were identified with sequential numbers. The first recording was taken soon after flag leaf emergence, the second was taken again after anthesis, and the third in the early milk stage. The trials with only one observation were recorded right after anthesis for responses at the adult plant stage as stripe rust was not present in the field duringearlier growth stages. IT was recorded based on a 0-9 scale (Line and Qayoum, 1992). SEV was recorded as a percentage of the leaf infected area using a modified Cobb Scale (Peterson et al., 1948). Table 1 summarizes environments, years, genotyped individuals, and measurements taken for each trial where stripe rust was recorded.

In order to account for differences in disease pressure in different environments, a two-step adjusted mean method was used in which a linear model was implemented to adjust both IT and SEV means within and across environments. Then a mixed linear model was used to calculate genomic estimated breeding values (GEBVs; Ward et al., 2019). Adjusted means from the stripe rust data collected in the unreplicated trials were adjusted using residuals calculated for the unreplicated genotypes in individual environments and across environments using the modified augmented complete block design model (ACBD; Federer 1956; Goldman 2019). The adjustments were made following the method implemented in Merrick and Carter (2021), with the full model across environments as follows:

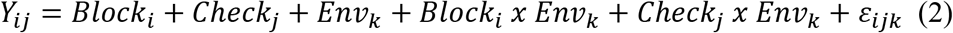

where *Y*_*ij*_ is the trait of interest; *Block*_*i*_ is the fixed effect of the ith block; *Check*_*j*_ is the fixed effect of the jth replicated check cultivar; *Env*_*k*_ is the fixed effect of the kth environment; and *ε*_*ijk*_ are the residual errors.

Heritability on a genotype-difference basis for broad-sense heritability was calculated using the variance components from the models implemented in Merrick and Carter (Merrick and Carter, 2021) and using best linear unbiased predictors for both individual environments and across environments using the formula:

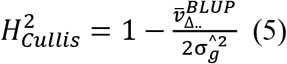

where 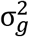 and *V̅*BLUP are the genotype variance and mean-variance of the BLUPs, respectively (Cullis et al., 2006).

### Genotypic Data

Lines were genotyped using genotyped-by-sequencing (GBS; Elshire et al., 2011) through the North Carolina State Genomics Sciences Laboratory in Raleigh, North Carolina, using the restriction enzymes *Msp*I and *Pst*I (Poland et al., 2012). Genomic DNA was isolated from seedlings in the one to three leaf stage using Qiagen BioSprint 96 Plant kits and the Qiagen BioSprint 96 workstation (Qiagen, Germantown, MD). DNA libraries were prepared following the protocol of DNA digestion with *Pst*I and *Msp*I restriction enzymes (Poland et al, 2012). Genotyping by sequencing (GBS; (Elshire et al., 2011) was conducted at North Carolina State University Genomic Sciences Laboratory with either an Illumina HiSeq 2500 or a NovaSeq 6000. DNA library barcode adapters, DNA library analysis, and sequence SNP calling were provided by the USDA Eastern Regional Small Grains Genotyping Laboratory (Raleigh, NC). Sequences were aligned to the Chinese Spring International Wheat Genome Sequencing Consortium (IWGSC) RefSeq v1.0 (Appels et al., 2018), using the Burrows-Wheeler Aligner (BWA) 0.7.17 (Li and Durbin, 2009). Genetic markers with more than 20% missing data, minor allele frequency of less than 5%, and those that were monomorphic were removed. Markers were then imputed using Beagle version 5.0 and filtered once more for markers under a 5% minor allele frequency (Browning et al., 2018). A total of 31,975 single-nucleotide polymorphism (SNP) markers for the 475 unique DP lines and the 2,630 BL lines were obtained from GBS. Principal components for the markers were calculated using the function ‘prcomp’, and a biplot with k-mean clusters was creating using the function “autoplot” in R (R Core Team, 2018).

Major rust resistance genes observed to be common in the WSU breeding population are for *Yr10*, *Yr17*, *Lr68*, and *Qyr.wpg-1B.1*and molecular marker data for these genes were included as fixed effects in our GS models. All winter wheat lines were genotyped with Kompetitive Allele Specific PCR (KASP®) assay for *Yr17*, *Lr68*, and *Qyr.wpg-1B.1* in the WSU winter wheat breeding laboratory. The *Yr17* gene (Helguera et al., 2003) was screened using the KASP marker developed by Milus, Lee, and Brown-Guedira (2015). The *Lr68* leaf rust resistance gene (Herrera-Foessel et al., 2012) was screened using the KASP marker developed by Rasheed et al. (2016). Although leaf rust resistance is not commonly selected for in the US Pacific Northwest (PNW) breeding programs, this gene was found in a large proportion of breeding lines, and thus was hypothesized that it may have been selected congruently with stripe rust resistance. The APR QTL *Qyr.wpg-1B.1*reported on chromosome 1B by Naruoka et al. (2015) was screened using the marker *IWB12603* (Mu et al., 2020). The KASP assays were performed using PACE^TM^ Genotyping Master Mix (3CR Bioscience, Essex, UK) following the manufacturer’s instructions and endpoint genotyping was conducted from fluorescence using a Lightcycler 480 Instrument II (Roche, Indianpolis, IN). The previously reported ASR gene *Yr10* (Frick et al., 1998) was screened with microsatellite marker *Xpsp3000* developed by Bariana et al. (2002). The microsatellite marker *Xpsp3000* was run using polymerase chain reaction (PCR) products and were separated on an ABI3730XL DNA fragment analyzer (Applied Biosystems), and alleles were scored with GeneMarkerv4.0 software (SoftGenetics), in collaboration with the USDA Western Regional Small Grains Genotyping Laboratory in Pullman, Washington.

### Genome-Wide Association Model

In addition to the inclusion of molecular markers for major rust resistance genes as fixed effects, markers identified through genome-wide association studies (GWAS) were included through de novo GWAS. This method is referred to further as genome-wide association study-assisted genomic selection (GWAS-GS). The GWAS- GS was implemented following McGowan et al. (2020). Briefly, GWAS was conducted using the Genome Association and Prediction Integrated Tool (GAPIT; Liu et al. 2016; Tang et al. 2016; Huang et al. 2019) with three principal components fitted as fixed effects on the training population or training fold. Three principal components were used because it was previously observed to be the most reliable in accounting for population structure for yield and agronomic traits in winter wheat for the same populations (Lozada et al., 2017). In accordance to advice put forward by Rice and Lipka (2019), the first method of GWAS-GS included only significant markers based on a Bonferonni cutoff of 0.05 (GWAS_B). For the remaining GWAS-GS methods, the markers were ordered by degree of statistical significance based on *p*-values from smallest to largest. We compared the inclusion of the top 5, 10, 25, 50, and 100 most significant markers as fixed effects (GWAS_5, GWAS_10, GWAS_25, GWAS_50, and GWAS_100).

### Prediction Models

#### Marker-Assisted Selection Model

Single and multiple regression models were used as MAS models to compare the major rust resistant markers and de novo GWAS markers’ predictive ability alone and in combination. The fixed effect multiple regression model is described as follows:

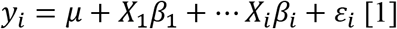

where *y*_*i*_ is the observed phenotypic value of the ith individual, *μ* is the mean, *X*_*i*_ is the genotype of marker i, and *β*_*i*_ is the effect of the ith marker, and *ε*_*i*_ is the residual error term.

#### Genomic Selection Model

rrBLUP was used as the base GS model and was implemented using the package ‘rrBLUP’ (Endelman, 2011). The basic rrBLUP model is described as follows (Rice and Lipka, 2019):

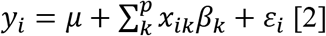

where *y*_*i*_ is the observed phenotypic value of the ith individual, *μ* is the mean, *x*_*ik*_ is the genotype of the kth marker and ith individual, *p* is the total number of markers, *β*_*k*_ is the estimated random marker effect of the kth marker, and *ε*_*i*_ is the residual error term.

#### Genomic Selection Model with Fixed Effects

To evaluate the effect of major and de novo GWAS markers on the prediction accuracy of the GS models, we used the rrBLUP model as described (Rice and Lipka, 2019):

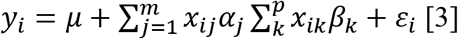

where *y*_*i*_ is the observed phenotypic value of the ith individual, *μ* is the mean, *x*_*ij*_ is the genotype of the jth individual, m is the number of markers included as fixed effect covariates, *α*_*j*_ is the fixed effect of the jth marker, *x*_*ik*_ is the genotype of the kth marker and ith individual, p is the total number of markers, *β*_*k*_ is the estimated random marker effect of the kth marker, and *ε*_*i*_ is the residual error term. The ridge regression penalty implemented by rrBLUP is not placed on the fixed effects, allowing a large effect on the model.

### Prediction Accuracy and Schemes

Prediction accuracy for the GS was reported using Pearson correlation coefficients between GEBVs and their respective adjusted means using the function “cor” in R (R Core Team, 2018). The effect of fixed-effect markers on prediction accuracy was assessed using a five-fold cross-validation scheme and independent validation sets for IT and SEV in the DP and BL training populations. The two populations were used to compare the effects of the significant markers in populations with different genetic relatedness, frequency of markers, and sources of resistant pedigrees. GS models were conducted with five-fold cross-validation by including 80% of the samples in the training population and predicting the GEBVs of the remaining 20% (Lozada and Carter, 2019). One replicate consists of the five model iterations, where the population is split into five different groups. This was completed 50 times. As mentioned previously for the GWAS-GS, the GWAS was conducted on 80% of the lines and then the markers are included in the GS model to predict the remaining 20% of the lines. Independent validation sets were then performed on a yearly basis by combining the two training populations and environments together per year. This allows the evaluation of models in a realistic breeding situation in which we combine all available data to build a training population.

The training populations were evaluated for cross-validations on a yearly basis and over combined years and trials. We assessed each year independently using cross-validations. We then created prediction models starting with the earliest trial and then a new model with the addition of each subsequent trial to evaluate genotype-by-environment interaction, continuous training of a prediction model, and the effect of different races of *P. striiformis* f. sp. *tritici*. The independent validation sets were first conducted using continuous training. For example, the earliest year i.e., 2013, was used to predict the following year i.e., 2014. The years were then combined to predict the following year i.e. 2013 and 2014 to predict 2015, and this process was continued until the years 2013 to 2018 were used to predict 2020.

All GS and MAS models and scenarios were analyzed using WSU’s Kamiak high performance computing cluster (Kamiak, 2021). Model comparisons were evaluated by using a Tukey’s honestly significant difference (HSD) test implemented in the “agricolae” package in R (R Core Team, 2018; de Mendiburu and de Mendiburu, 2019). The comparison of models was then plotted for visual comparison using “ggplot2” in R (Wickham, 2011; R Core Team, 2018).

## RESULTS

### Phenotypic Data

Stripe rust phenotyping was dependent on natural infection. Therefore, it is important to evaluate GS models in different years to account for environments with little to no variation in stripe rust severity and pathogen race changes. Overall, the maximum IT and SEV were relatively high for each scale, indicating the presence of adequate stripe rust severity for the checks present in each trial (Table 2). The BL had relatively high coefficient of variations (CV) for each trial. However, the heritability was very high, ranging from 0.60 to 0.96 across traits and trials, indicating adequate screening trials for stripe rust.

**Table 2.** Stripe rust infection type (IT) and disease severity (SEV) heritability (H^2^) and trial statistics for the diversity panel (DP and breeding line (BL) training population phenotypes from 2013 to 2016 and 2016 to 2020.

| Population | Year | Trait | $H^2$ | CV<sup>a</sup> | Max<sup>b</sup> | Mean | Min<sup>c</sup> | SD<sup>d</sup> |
| --- | --- | --- | --- | --- | --- | --- | --- | --- |
| DP | 2013 | IT | 0.85 | 52.31 | 9 | 3 | 1 | 2 |
| DP | 2014 | IT | 0.82 | 57.13 | 9 | 4 | 1 | 2 |
| DP | 2015 | IT | 0.89 | 43.82 | 9 | 5 | 1 | 2 |
| DP | 2016 | IT | 0.84 | 46.23 | 9 | 4 | 0 | 2 |
| DP | 2013-2014 | IT | 0.93 | 55.58 | 9 | 4 | 1 | 2 |
| DP | 2013-2015 | IT | 0.94 | 52.66 | 9 | 4 | 1 | 2 |
| DP | 2013-2016 | IT | 0.95 | 51.77 | 9 | 4 | 0 | 2 |
| DP | 2013 | SEV | 0.91 | 108.31 | 100 | 22 | 2 | 24 |
| DP | 2014 | SEV | 0.78 | 116.04 | 90 | 24 | 2 | 28 |
| DP | 2015 | SEV | 0.92 | 72.78 | 100 | 43 | 2 | 32 |
| DP | 2016 | SEV | 0.89 | 70.57 | 100 | 36 | 0 | 26 |
| DP | 2013-2014 | SEV | 0.93 | 112.77 | 100 | 23 | 2 | 26 |
| DP | 2013-2015 | SEV | 0.96 | 99.31 | 100 | 30 | 2 | 30 |
| DP | 2013-2016 | SEV | 0.96 | 94.90 | 100 | 31 | 0 | 29 |
| BL | 2016 | IT | 0.90 | 87.56 | 8 | 3 | 0 | 2 |
| BL | 2017 | IT | 0.83 | 83.36 | 9 | 3 | 0 | 2 |
| BL | 2018 | IT | 0.79 | 172.49 | 8 | 2 | 0 | 3 |
| BL | 2020 | IT | 0.96 | 93.03 | 8 | 3 | 0 | 3 |
| BL | 2016-2017 | IT | 0.83 | 84.06 | 9 | 3 | 0 | 2 |
| BL | 2016-2018 | IT | 0.79 | 115.26 | 9 | 2 | 0 | 3 |
| BL | 2016-2020 | IT | 0.80 | 113.22 | 9 | 2 | 0 | 3 |
| BL | 2016 | SEV | 0.86 | 152.31 | 80 | 9 | 0 | 13 |
| BL | 2017 | SEV | 0.88 | 131.79 | 90 | 16 | 0 | 21 |
| BL | 2018 | SEV | 0.60 | 212.74 | 80 | 11 | 0 | 24 |
| BL | 2020 | SEV | 0.96 | 125.06 | 80 | 18 | 0 | 23 |
| BL | 2016-2017 | SEV | 0.89 | 136.36 | 90 | 15 | 0 | 20 |
| BL | 2016-2018 | SEV | 0.85 | 165.25 | 90 | 13 | 0 | 22 |
| BL | 2016-2020 | SEV | 0.86 | 160.47 | 90 | 14 | 0 | 22 |

<sup>a</sup>CV: coefficient of variation
<sup>b</sup>Max: maximum
<sup>c</sup>Min: minimum
<sup>d</sup>SD: standard deviation

The inclusion of multiple environments creates a challenge for GS models due to the genotype-environment interaction (GEI). There were significant differences between each year for each population and trait (Figure 1A-D). The ranges for both IT and SEV were large, indicating both resistant and non-resistant varieties within the populations. The mean IT and SEV were also lower in the BL compared to the DP (Figure 1A-D; Table 2). The BL population consisted of a larger proportion of resistant cultivars compared to the DP, which was expected as these had previously been selected under field conditions. SEV displayed a large concentration of values near zero, specifically in the year 2018 (Figure 1D). Each year’s significant differences indicate an environmental effect that needs to be accounted for within prediction models.

**Figure 1.**
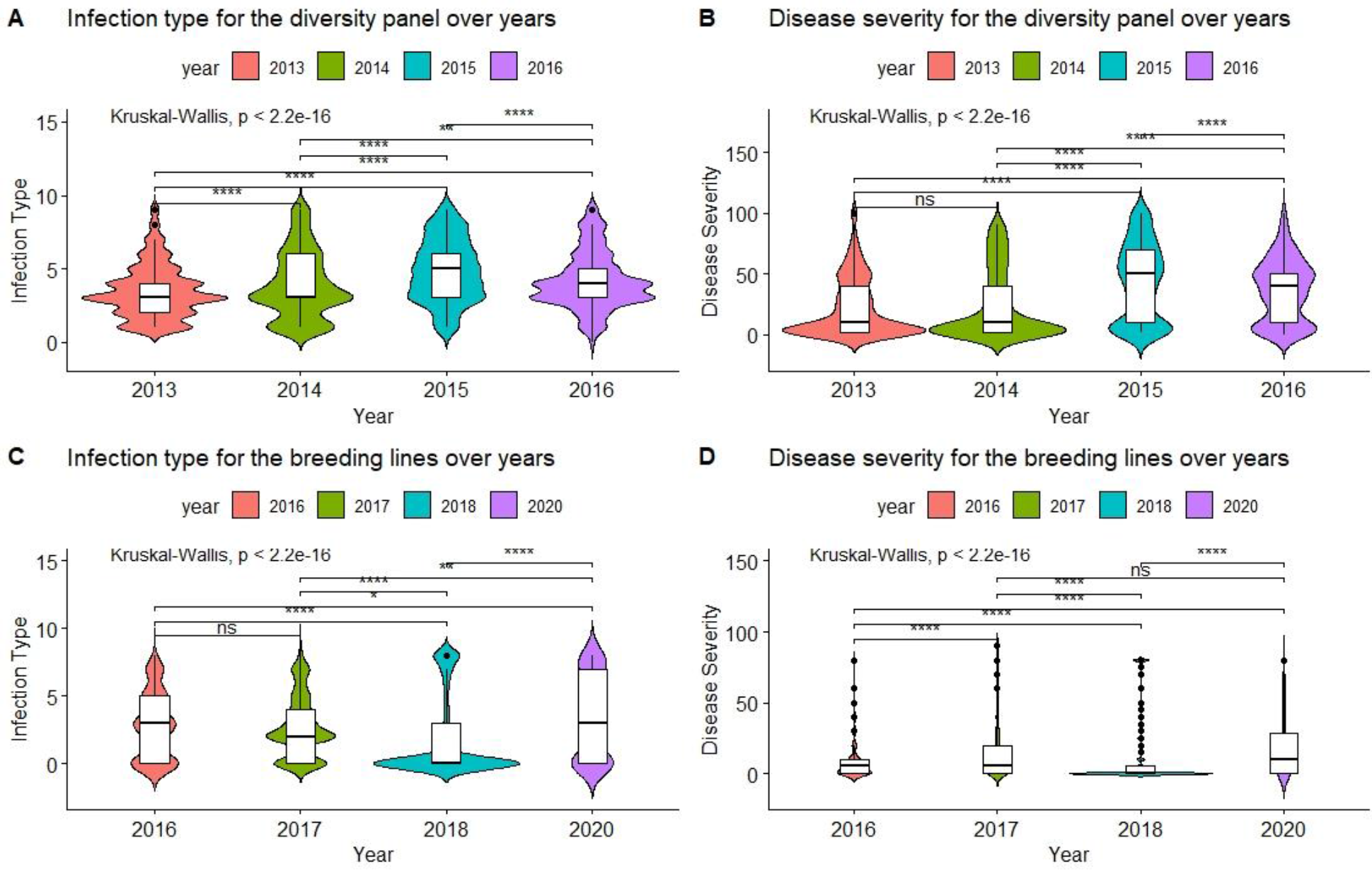
Comparison of infection type and disease severity over years in the diversity panel and breeding line training populations using Kruskal-Wallis H test.

In addition to GEI, stripe rust races may change from year to year, which creates the opportunity for major genes to be overcome by virulent races. The USDA stripe rust lab records race frequencies each year. The major stripe rust races for each year was either PSTv-37 or PSTv-52 with the exception of 2017 and 2020, which had large frequencies for PSTv-37 (Table S1). The other races with higher frequencies included PSTv-39, PSTv-322, PSTv-48, PSTv-79, PSTv-11, and PSTv-73. Therefore, the difference in race change was not a major factor in prediction scenarios.

### Genotypic Data

The major rust genes present within the WSU winter wheat breeding program germplasm are for *Yr10*, *Yr17*, *Lr68*, and *Qyr.wpg-1B.* The frequency of genotypes, as determined by the previously described molecular markers for each of these genes, are presented in Table 3. Similar frequencies in both populations was observed for the homozygous resistant allele for *Lr68* with 50% and 46% in the DP and BL, respectively. The frequency of the marker for *Yr10* and *IWB12603* were much higher in the DP than in the BL with *Yr10* having a relatively high frequency of 53% in the DP. However, the homozygous resistant allele for *Yr17* was much higher in the BL (38%) than in the DP (19%). There was also a wide combination of homozygous resistant alleles within each population (Figure 2).

**Table 3.** Frequency of rust resistant genotypes present in both the breeding line (BL) and diversity panel line (DP) populations.

| Population | Marker (gene) | Genotype | Number<sup>b</sup> | Frequency |
| --- | --- | --- | --- | --- |
| DP | KASP (*Yr17*) | 0 | 356 | 0.75 |
|  |  | 1 | 30 | 0.06 |
|  |  | 2 | 89 | 0.19 |
| DP | IWB12603(Qyr.wpg-1B.1) | 0 | 288 | 0.61 |
|  |  | 1 | 16 | 0.03 |
|  |  | 2 | 171 | 0.36 |
| DP | KASP (*Lr68*) | 0 | 182 | 0.38 |
|  |  | 1 | 55 | 0.12 |
|  |  | 2 | 238 | 0.50 |
| DP | Xpsp3000(*Yr10*) | 0 | 220 | 0.46 |
|  |  | 1 | 4 | 0.01 |
|  |  | 2 | 251 | 0.53 |
| BL | KASP(*Yr17*) | 0 | 1491 | 0.57 |
|  |  | 1 | 131 | 0.05 |
|  |  | 2 | 1008 | 0.38 |
| BL | IWB12603(Qyr.wpg-1B.1) | 0 | 2244 | 0.85 |
|  |  | 1 | 53 | 0.02 |
|  |  | 2 | 333 | 0.13 |
| BL | KASP(*Lr68*) | 0 | 1255 | 0.48 |
|  |  | 1 | 166 | 0.06 |
|  |  | 2 | 1209 | 0.46 |
| BL | Xpsp3000 (*Yr10*) | 0 | 2172 | 0.83 |
|  |  | 1 | 11 | 0.00 |
|  |  | 2 | 447 | 0.17 |

<sup>a</sup>Genotype: Allele 0: homozygous wild type allele; Allele 1: heterozygous with both alleles present; Allele 2: homozygous resistant allele
<sup>b</sup>Number: number of lines.

**Figure 2.**
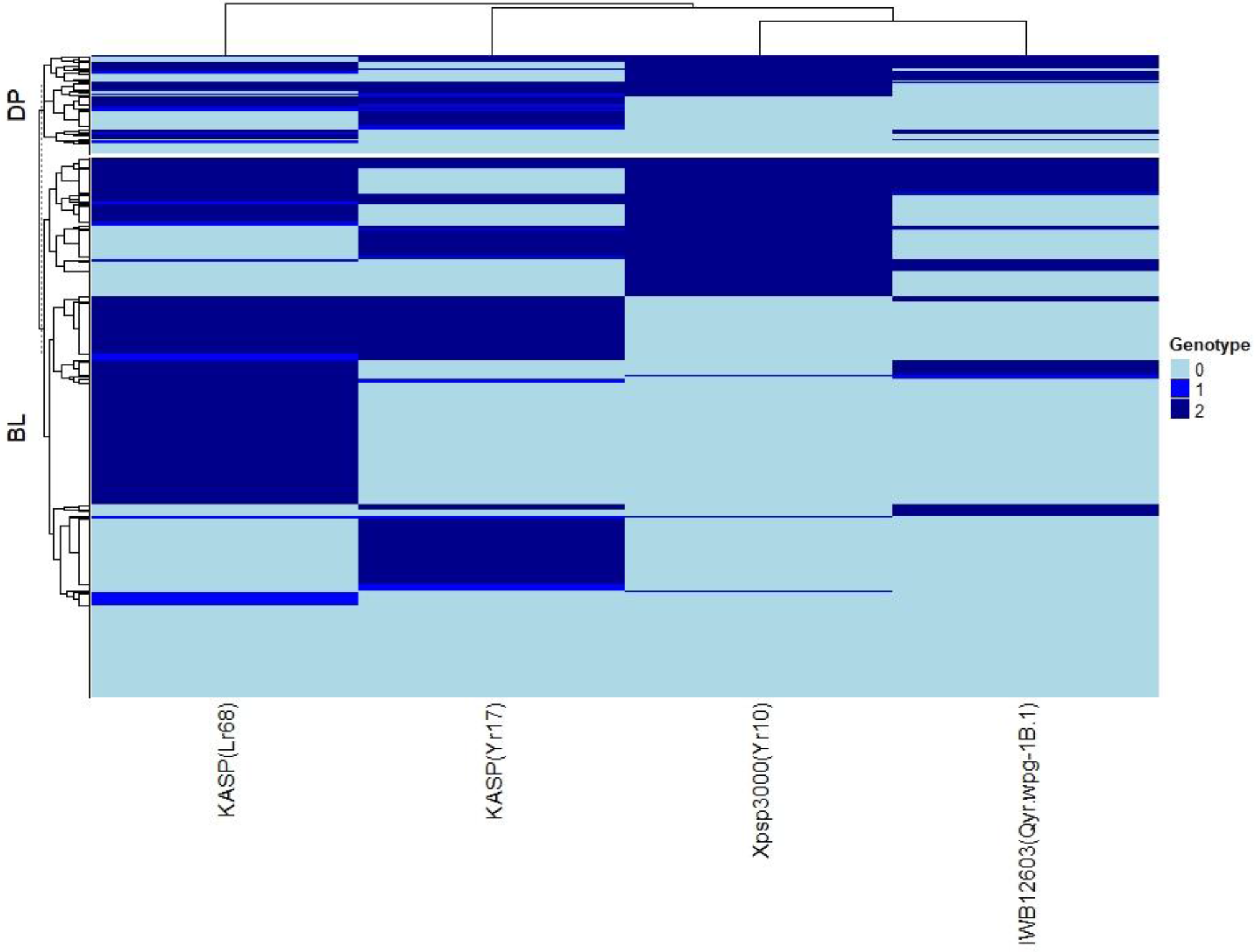
Heat map and hierarchical clustering for lines in the diversity panel (DP) and breeding line (BL) populations for major rust markers: IWB12603(Qyr.wpg-1B.1), KASP(Lr68), Xpsp3000(Yr10), and KASP(Yr17). Genotype: 0: homozygous wild type allele; 1: heterozygous with both alleles present; 2: homozygous resistant allele

The principal component biplot using the GBS SNP markers over the combined DP and BL training populations accounted for only 9.1% of the total genetic variation, indicating a large population structure (Figure 3). PC1 explained 5.4% of the variation, and PC2 explained 3.7% of the variation. The biplot revealed three main clusters over the combined populations using k-means clustering.

**Figure 3.**
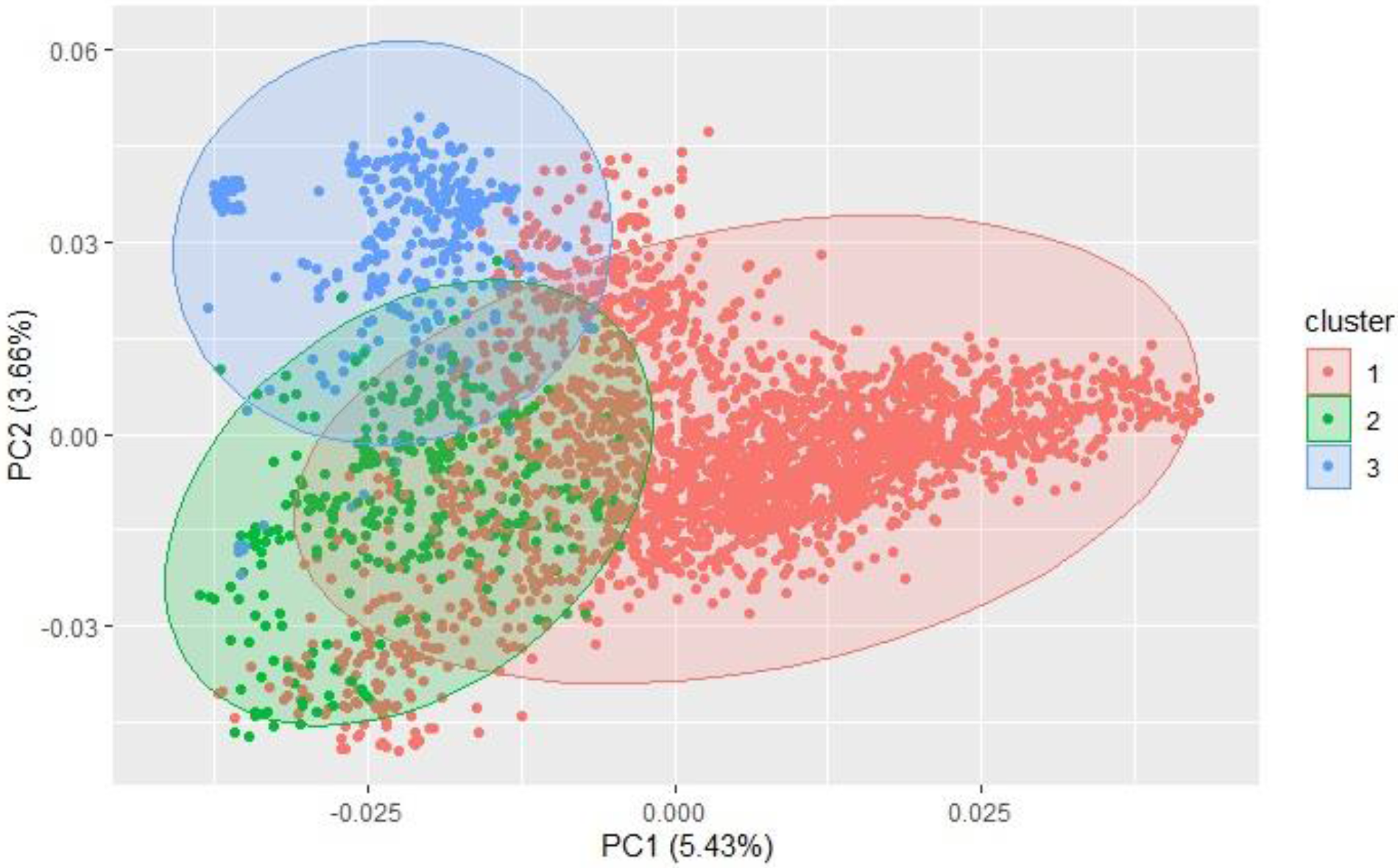
Principal component (PC) biplot and kmeans clustering of SNP GBS markers from the diversity panel and breeding line training populations.

### Cross-Validations

#### Major Markers

Multiple comparisons between the inclusion of each major molecular marker for known rust resistance genes individually and in combination (ALL_M) were completed for both populations in and across years for IT and SEV (Table S2 and S3). The markers for major rust genes were included as fixed effects and compared to the base rrBLUP model and MAS models with the markers as variables alone (Table S4 and S5). Within individual years in the BL, the rrBLUP base model reached a high accuracy of 0.65 within 2018 and 2020 for IT and 0.68 within 2018 for SEV. The major markers’ effects varied from year to year, but the marker for *Yr17* showed an increase in prediction accuracy for every year except in 2018 for IT (Table S2) and in 2018 and across 2016-2018 and 2016-2020 for SEV (Table S3). The majority of markers had relatively low prediction accuracies for MAS with the exception of the *Yr17* marker that reached an accuracy of .05 and 0.42 for IT and SEV, respectively (Table S4 and S5). When all markers were combined, similar accuracies were reached compared to when only the *Yr17* marker was included in both the rrBLUP and MAS models. The remainder of the major rust gene markers with the exception of *Lr68* increased accuracy within specific years but had less consistency than the marker for *Yr17*. The largest differences from the rrBLUP model within a single year in the BL were seen in 2016 for the GS models (Figure 4). Within 2016, the combination of both *Yr10* and *Yr17* markers increased accuracy by 0.06 for SEV. *Yr17* and the combination of markers only slightly increased accuracy across environments with an increase in IT of 0.01 (Figure 5).

**Figure 4.**
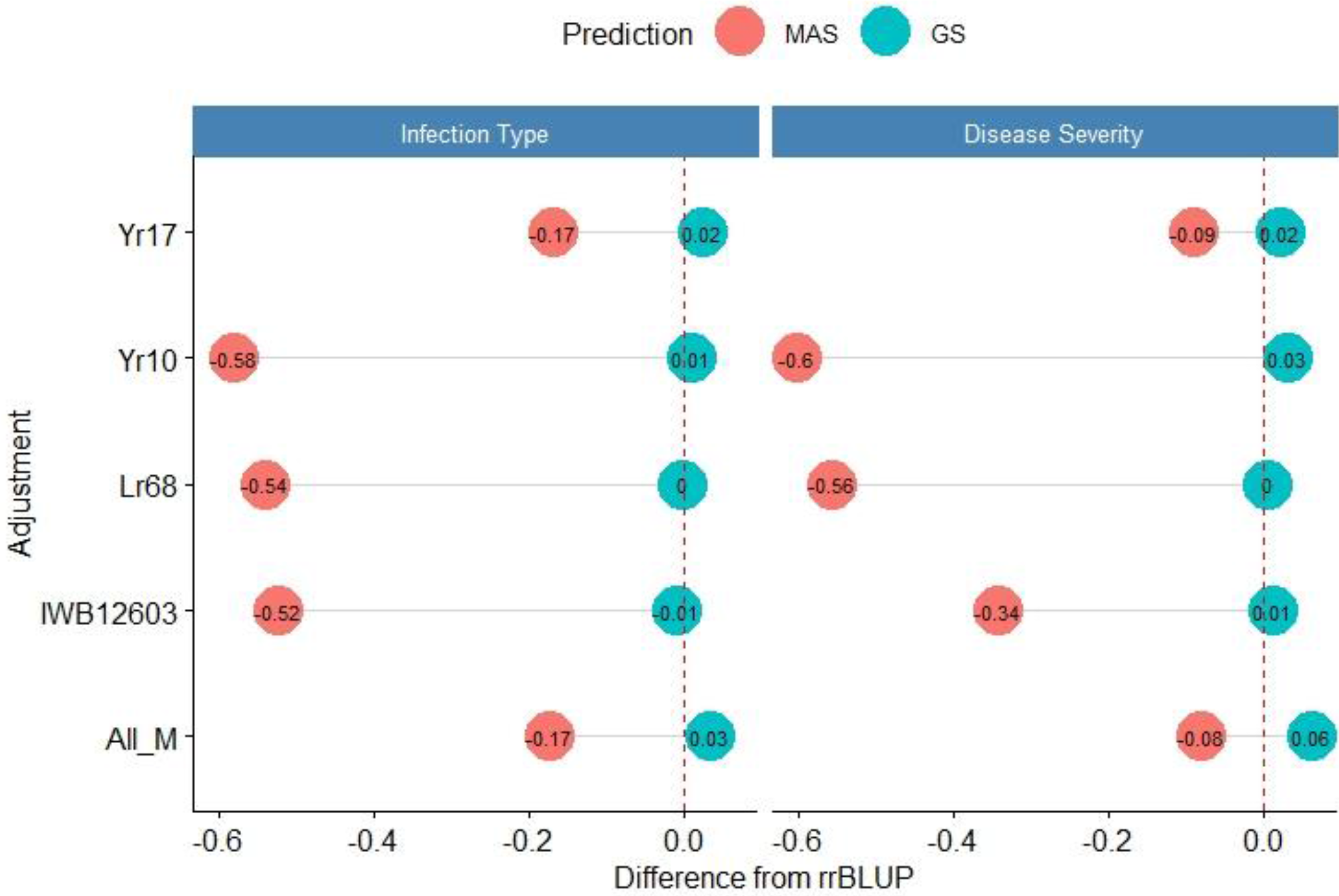
Difference from the base rrBLUP model for major markers in genomic selection (GS) and marker-assisted selection (MAS) in the breeding lines in 2016. Adjustments: ALL_M: IWB12603(Qyr.wpg-1B.1), KASP(Lr68), Xpsp3000(Yr10), and KASP(Yr17) combined.

**Figure 5.**
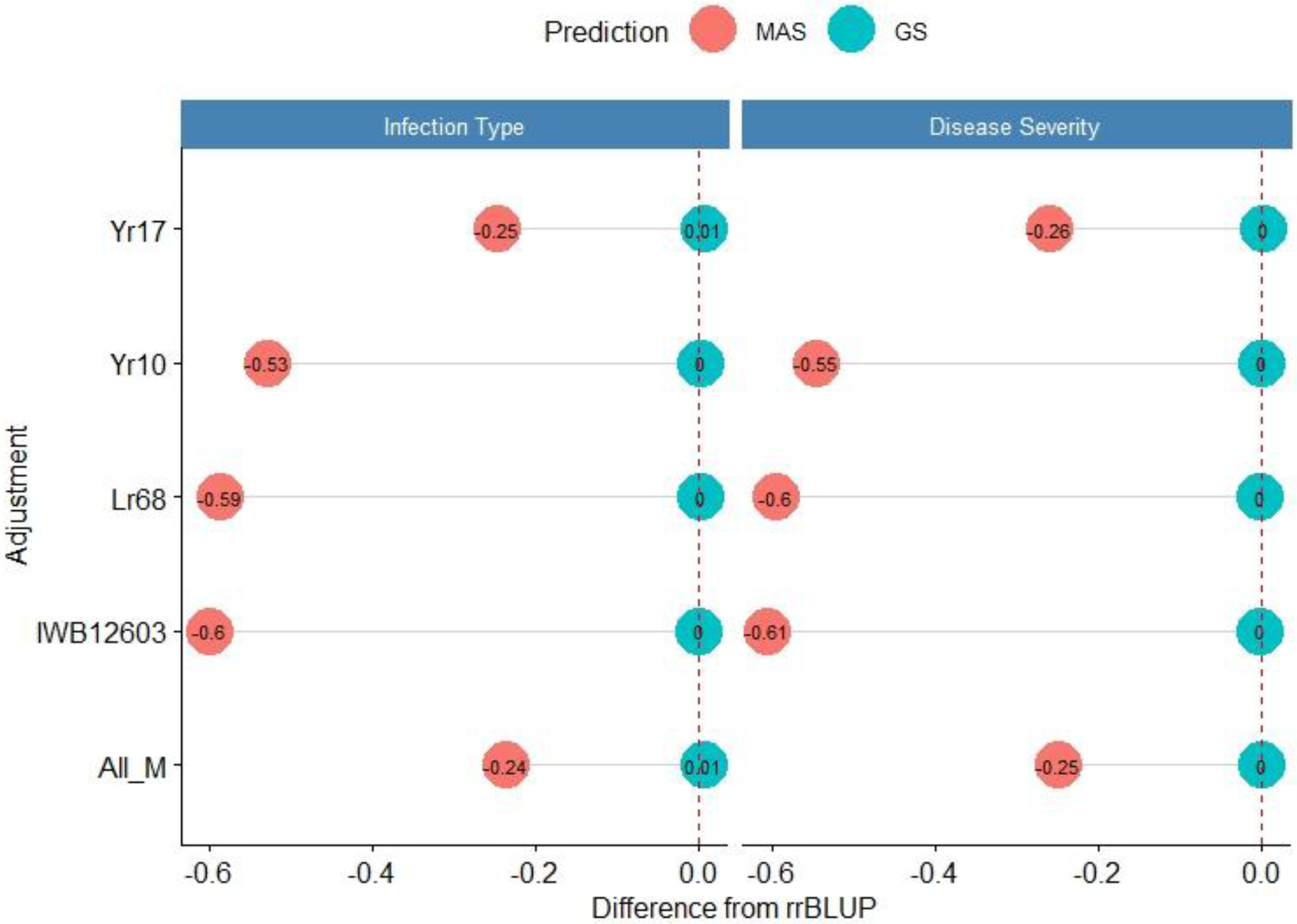
Difference from the base rrBLUP model for major markers genomic selection (GS) and marker-assisted selection (MAS) in the breeding lines across 2016-2020. Adjustments: ALL_M: IWB12603(Qyr.wpg-1B.1), KASP(Lr68), Xpsp3000(Yr10), and KASP(Yr17) combined.

Within individual years in the DP, the rrBLUP base model reached an accuracy of 0.55 for IT (Table S2) and 0.64 for SEV in 2013 (Table S3). Across years, IT reached 0.56 in 2013-2016 (Table S2) and 0.69 for SEV in 2013-2014 (Table S3). Within the DP, the major rust markers had less of an effect on prediction accuracy, with *Yr10* being the only marker that increased accuracy from the base rrBLUP model and at a maximum of 0.01. For MAS, the combination of markers resulted in the least reduction of accuracy with a maximum reduction of 0.10 within 2015 for IT and 0.07 within 2016 (Table S4) for SEV (Table S5). Markers for *Yr10* and IWB12603 also had the largest effect on the MAS models. The largest differences from the rrBLUP model within a single year in the DP were seen in 2015 for the GS models (Figure 6). Within 2015, the *Yr10* marker increased accuracy by 0.01. There were no increases in accuracy across any combination of environments in the DP.

**Figure 6.**
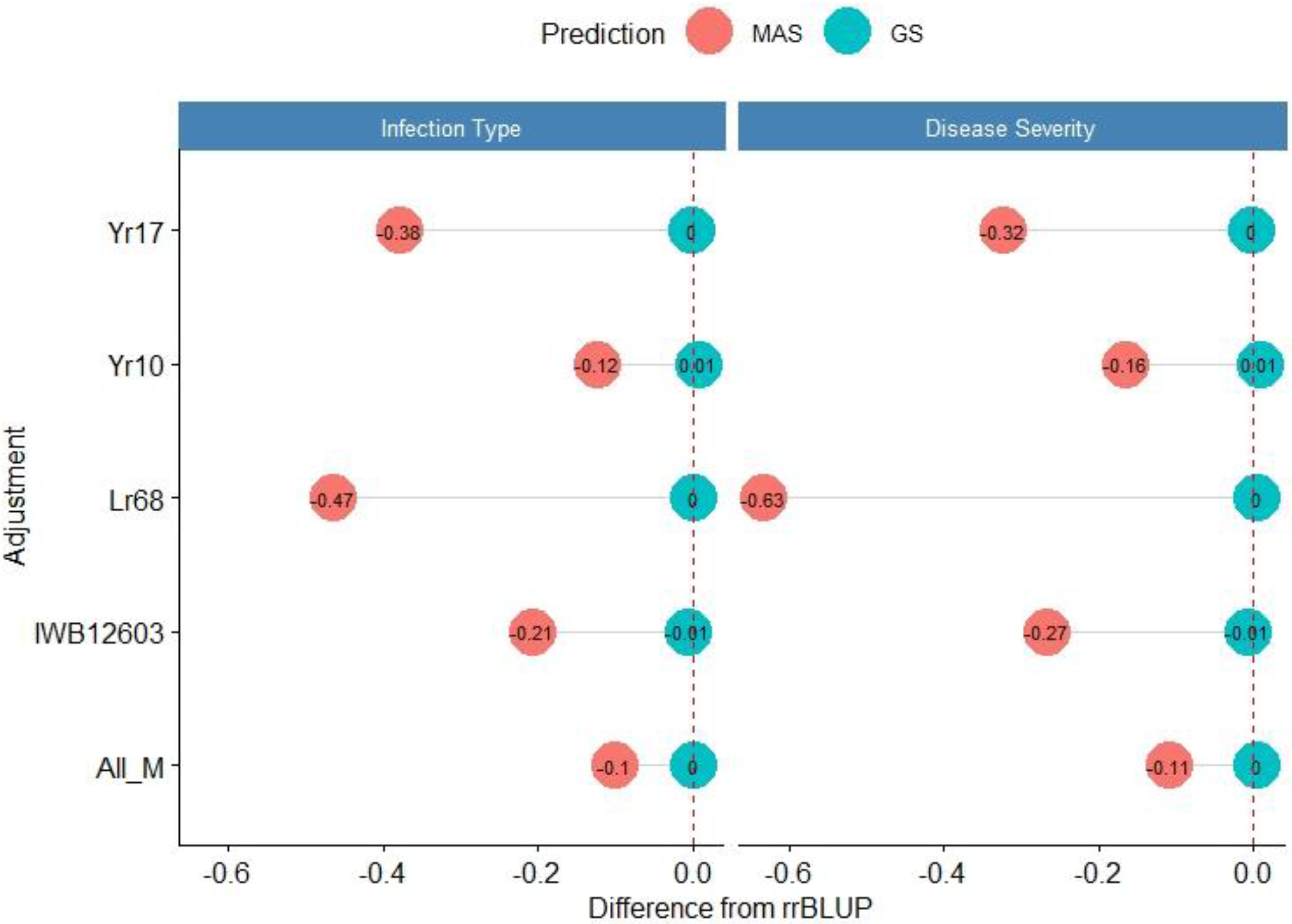
Difference from the base rrBLUP model for major markers in genomic selection (GS) and marker-assisted selection (MAS) in the diversity panel lines in 2015. Adjustments: ALL_M: IWB12603(Qyr.wpg-1B.1), KASP(Lr68), Xpsp3000(Yr10), and KASP(Yr17) combined.

#### De Novo GWAS Markers

The de novo GWAS markers increased prediction accuracy in individual years and across years in the BL, but not in the DP. Only the GWAS_B, GWAS_5, and GWAS_10 sets increased accuracy with GWAS_25, GWAS_50, and GWAS_100 decreasing prediction accuracy. The largest increase in IT was for GWAS_5 in 2018 with an increase of 0.02 for both IT and SEV (Table S1 and S2; Figure 7). Across years, GWAS_10 had the largest increase of 0.02 in 2016-2018 for SEV (Table S3; Figure 8). The MAS for the de novo GWAS markers had larger decreases in MAS compared to the major markers in both the DP and BL (Table S4 and S5). The larger GWAS sets (GWAS_25, GWAS_50, and GWAS_100) consistently had lower prediction accuracies than the other GWAS sets and the major rust gene markers. GWAS_B using significant markers showed similar accuracies to GWAS_5 displaying no advantage compared to arbitrarily including markers based on the *p*-value.

**Figure 7.**
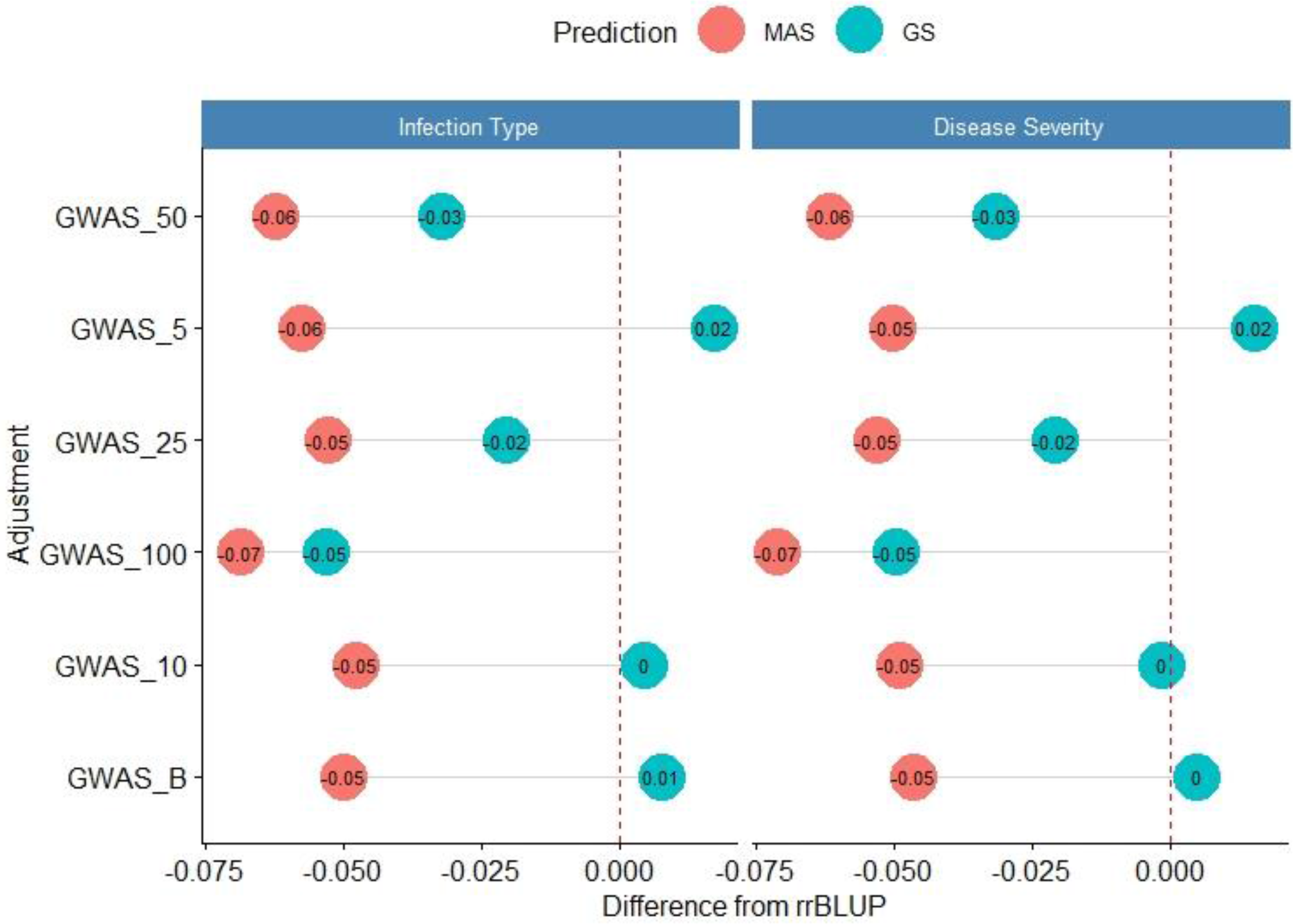
Difference from the base rrBLUP model for de novo GWAS markers in genomic selection (GS) and marker-assisted selection (MAS) in the breeding lines in 2018. Adjustments: GWAS_B: genome-wide association assisted genomic selection (GWAS-GS) with Bonferonni significant markers; GWAS_5: GWAS-GS with the top 5 significant markers; GWAS_10: GWAS-GS with the top 10 significant markers; GWAS_25: GWAS-GS with the top 25 significant markers; GWAS_50: GWAS-GS with the top 50 significant markers; and GWAS_100: GWAS-GS with the top 100 significant markers.

**Figure 8.**
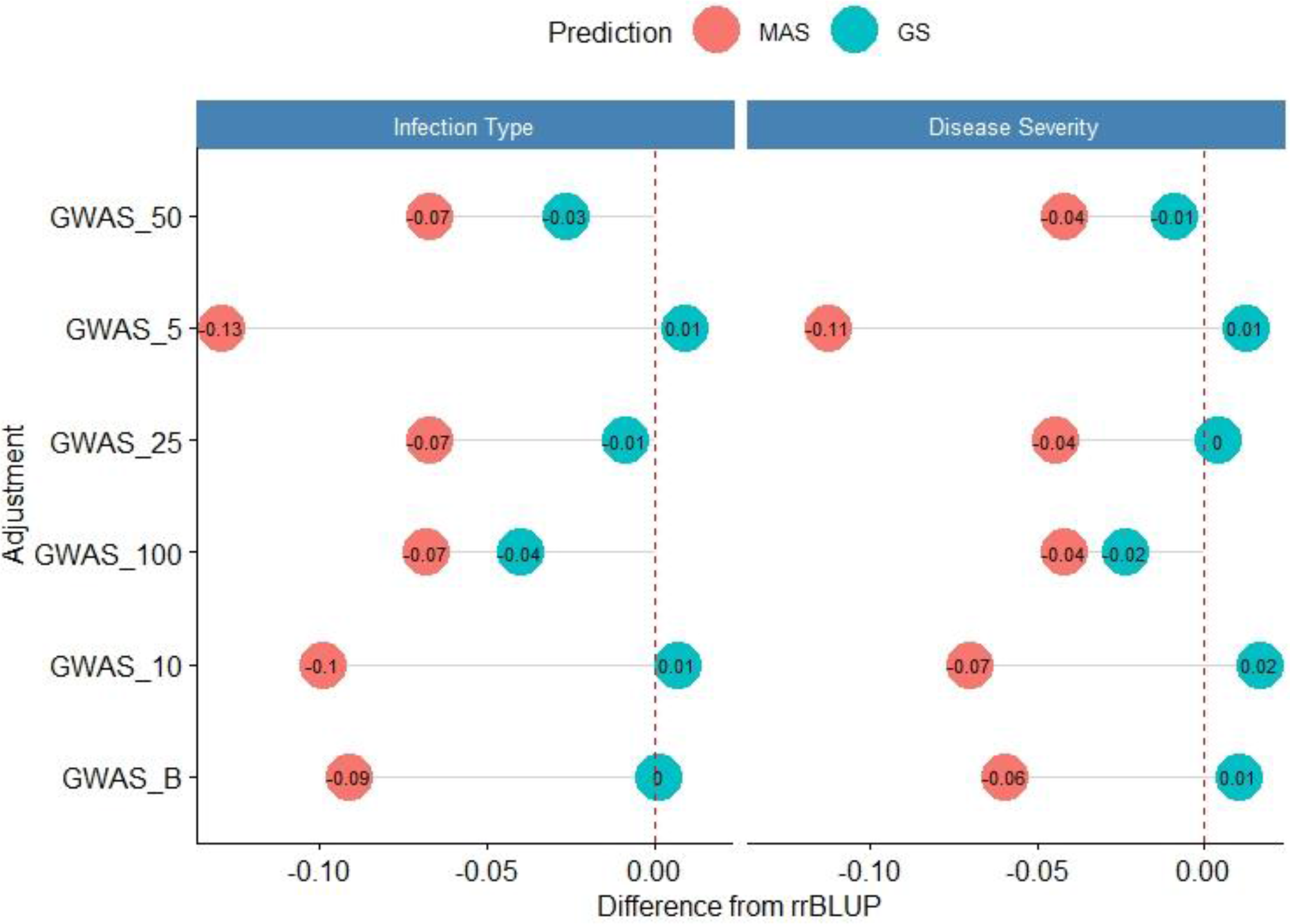
Difference from the base rrBLUP model for de novo GWAS markers in genomic selection (GS) and marker-assisted selection (MAS) in the breeding lines across 2016-2018. Adjustments: GWAS_B: genome-wide association assisted genomic selection (GWAS-GS) with Bonferonni significant markers; GWAS_5: GWAS-GS with the top 5 significant markers; GWAS_10: GWAS-GS with the top 10 significant markers; GWAS_25: GWAS-GS with the top 25 significant markers; GWAS_50: GWAS-GS with the top 50 significant markers; GWAS_100: GWAS-GS with the top 100 significant markers.

### Validation Sets

#### Major Markers

The validation sets were conducted by combining both training populations and years and predicting the following year as a forward prediction. In doing so, the validation sets were evaluated to demonstrate real-world breeding scenarios wherein all information available was used to create predictions. The first three years, 2013-2015, consisted exclusively of the DP, and from 2016 forward, the BL was included due to the availability of training populations. The validation sets resulted in the highest accuracy out of all prediction scenarios with the rrBLUP base model and all of the major markers reaching an accuracy of 0.72 in the SEV for predicting 2014 using the 2013 data (Table S6 and S7). For the same year, the major markers with the exception of *Yr10* resulted in an increase of accuracy of 0.01. The major rust markers either performed the same or increased accuracy for the majority of validation GS predictions.

As the number of environments and years were added to the population, the general prediction accuracy decreased presumably due to the prediction of multiple environments within a year and the inclusion of different training populations. However, as the accuracy decreased for the base rrBLUP model, the fixed markers’ effect increased. The largest increase in both the cross-validations and validation sets occurred using 2013-2017 to predict 2018, resulting in the MAS models using *Yr17* and all markers with an increase of 0.17 and 0.16 respectively for IT and 0.13 and 0.10 respectively for SEV (Table S8 and S9; Figure 9). The validation sets were the only prediction scenarios in which MAS performed better than GS models. However, this was not the case for all MAS models, with most major markers showing similar decreases in accuracy compared to cross-validations.

**Figure 9.**
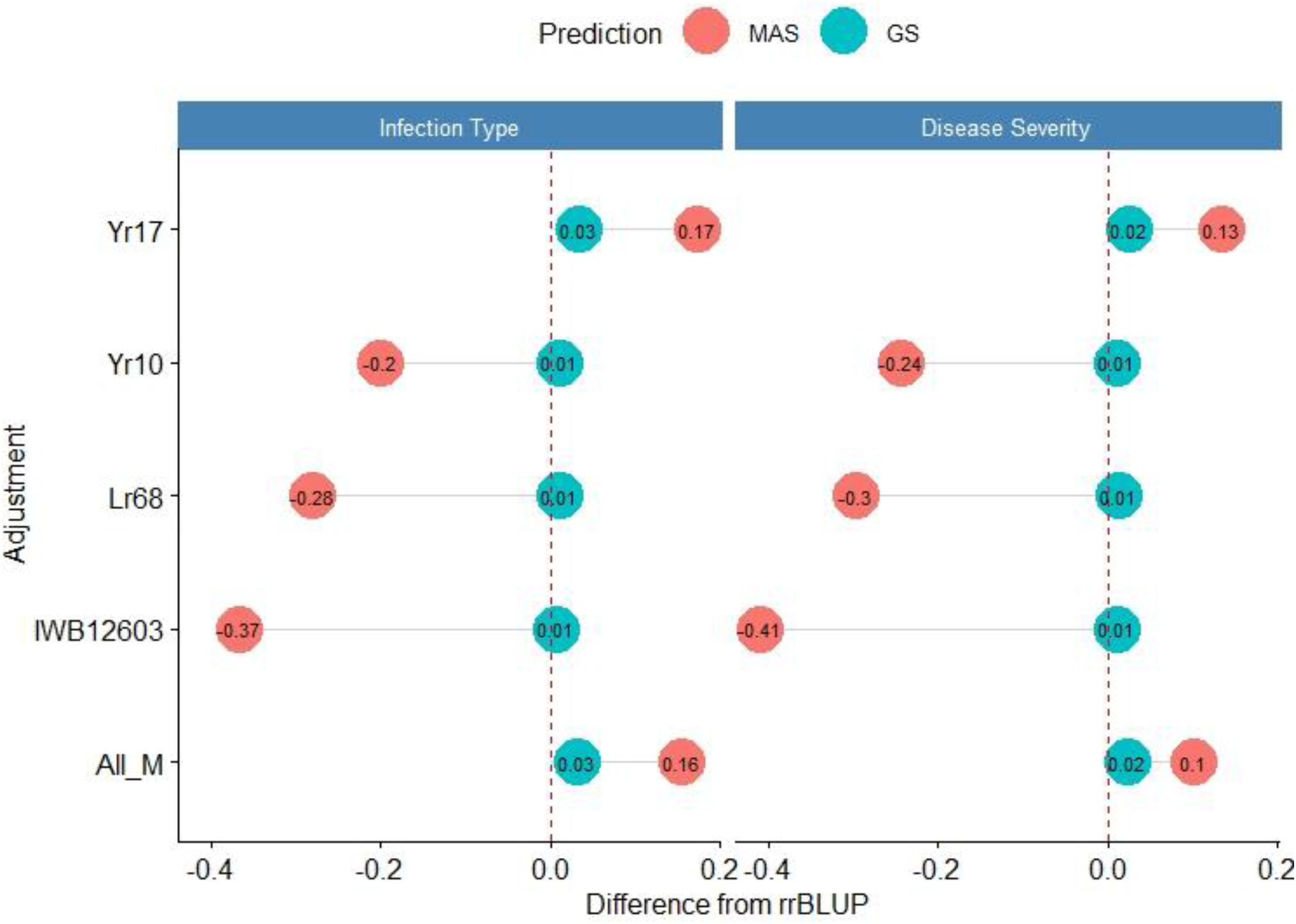
Difference from the base rrBLUP model for major markers in genomic selection (GS) and marker-assisted selection (MAS)in the validation set using the years 2013-2017 to predict 2018. Adjustments: ALL_M: IWB12603(Qyr.wpg-1B.1), KASP(Lr68), Xpsp3000(Yr10), and KASP(Yr17) combined.

#### De Novo GWAS Markers

The de novo GWAS marker sets also increased accuracy when more environments were included. The increase of prediction accuracy was not seen in the earlier validation sets as seen for the molecular markers for major rust genes. The de novo GWAS markers had the largest prediction accuracies in the last two validation sets with GWAS_5 having an accuracy of 0.33 for IT and GWAS having an accuracy of 0.38 for SEV using 2013-2017 to predict 2018 (Table S6 and S7). In the last validation set, GWAS had the largest prediction accuracy of 0.55 for IT. Similarly, the smaller GWAS sets had the highest prediction accuracy. In contrast to the cross-validations, the larger GWAS sets did not have a drastic decrease with GWAS_100, and actually had the same prediction accuracy as the base rrBLUP for IT and an increase of 0.01 for SEV in using 2013-2018 to predict 2020 (Table S6 and S7). The de novo GWAS marker sets had the largest increases over all scenarios with the GWAS_5 having an increase of 0.19 with MAS for IT (Table S8 and S9; Figure 10).

**Figure 10.**
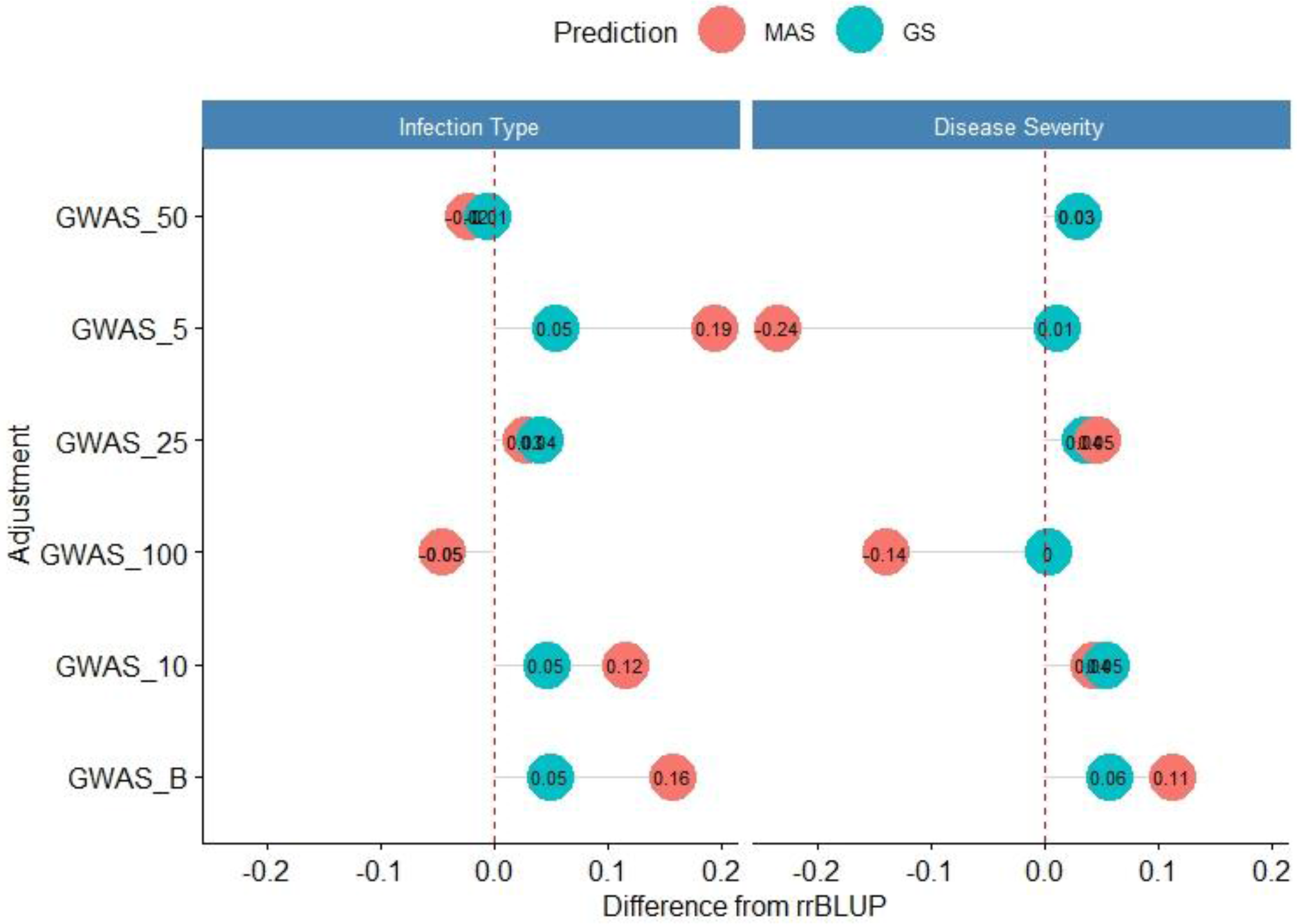
Difference from the base rrBLUP Model for de novo GWAS markers in genomic selection (GS) and marker-assisted selection (MAS)in the validation set using the years 2013-2017 to predict 2018. Adjustments: GWAS_B: genome-wide association assisted genomic selection (GWAS-GS) with Bonferonni significant markers; GWAS_5: GWAS-GS with the top 5 significant markers; GWAS_10: GWAS-GS with the top 10 significant markers; GWAS_25: GWAS-GS with the top 25 significant markers; GWAS_50: GWAS-GS with the top 50 significant markers; and GWAS_100: GWAS-GS with the top 100 significant markers.

### Overall Differences

When we compared the different models over all years within each population, we found that the marker for *Yr17* and the combination of all markers had the largest prediction accuracies. However, the increase was only statistically significant in the BL population and in the validation sets. There was no statistical increase in prediction accuracy in the DP. The largest mean accuracy in any population was the major rust markers and base rrBLUP for SEV in the DP with an accuracy of 0.64 across all years (Table 4). There was also a statistical increase in prediction accuracy as we increased the combination of years over both IT and SEV training populations with an accuracy of 0.57 and 0.63 for IT and SEV, respectively, when years 1-4 were combined (Table 5).

**Table 4.** Comparison of genomic selection models accuracy and pairwise comparisons for stripe rust infection type (IT) and disease severity (SEV) for Pacific Northwest winter wheat diversity panel (DP) lines and breeding lines (BL) phenotyped from 2013-2020 in Central Ferry, Lind, and Pullman, WA over all individual population cross-validation sets and combined validation sets.

| Trait | Pop | All_M | GWAS_B | GWAS_10 | GWAS_100 | GWAS_25 | GWAS_5 | GWAS_50 | IWB12603 | *Lr68* | rrBLUP | *Yr10* | *Yr17* |
| --- | --- | --- | --- | --- | --- | --- | --- | --- | --- | --- | --- | --- | --- |
| IT | BL | 0.59a | 0.57c | 0.56d | 0.51g | 0.54e | 0.57c | 0.52f | 0.58b | 0.58b | 0.58b | 0.58b | 0.59a |
|  | DP | 0.52b | 0.47c | 0.44d | 0.4g | 0.42e | 0.47c | 0.41f | 0.52b | 0.53ab | 0.53ab | 0.53a | 0.53ab |
|  | VS | 0.49ab | 0.5a | 0.48d | 0.43g | 0.45e | 0.49bc | 0.44f | 0.48d | 0.48d | 0.48d | 0.49cd | 0.49b |
| SEV | BL | 0.6a | 0.57cd | 0.56d | 0.52g | 0.55e | 0.58c | 0.53f | 0.59b | 0.59b | 0.59b | 0.59ab | 0.6ab |
|  | DP | 0.64ab | 0.6c | 0.57d | 0.53g | 0.54e | 0.6c | 0.54f | 0.63b | 0.64ab | 0.64ab | 0.64a | 0.64ab |
|  | VS | 0.55ab | 0.54cd | 0.54cd | 0.52e | 0.52e | 0.54d | 0.52e | 0.54abc | 0.54abc | 0.54bc | 0.54abc | 0.55a |

Models labeled with the same letter are not significantly different ( $P$ -value = 0.05). Adjustments: ALL\_M: IWB12603(Qyr.wpg-1B.1), KASP (*Lr68*), Xpsp3000(*Yr10*), and KASP (*Yr17*) combined; GWAS\_B: genome-wide association assisted genomic selection (GWAS-GS) with Bonferonni significant markers; GWAS\_5: GWAS-GS with the top 5 significant markers; GWAS\_10: GWAS-GS with the top 10 significant markers; GWAS\_25: GWAS-GS with the top 25 significant markers; GWAS\_50: GWAS-GS with the top 50 significant markers; and GWAS\_100: GWAS-GS with the top 100 significant markers.

**Table 5.** Comparison of the number of years in the training populations on overall genomic selection model accuracy and pairwise comparisons for stripe rust infection type (IT) and disease severity (SEV) for Pacific Northwest winter wheat over both the diversity panel lines and breeding lines phenotyped from 2013-2020 in Central Ferry, Lind, and Pullman, WA in the cross-validation sets.

| Trait | Year 1 | Year 2 | Year 3 | Year 4 | Year 1-2 | Year 1-3 | Year 1-4 |
| --- | --- | --- | --- | --- | --- | --- | --- |
| IT | 0.52d | 0.43f | 0.57a | 0.54c | 0.49e | 0.56b | 0.57a |
| SEV | 0.53f | 0.55e | 0.61b | 0.59c | 0.58d | 0.62a | 0.63a |

Models labeled with the same letter are not significantly different ( $P$ -value = 0.05).

## DISCUSSION

### Genomic Selection for Disease Resistance

The development of resistant cultivars is the most effective and economical method for controlling diseases such as stripe rust (Chen and Kang, 2017). Due to the challenges of breeding for both quantitative and qualitative disease resistance, it is recommended to combine them. In addition to the challenges for breeding both major gene qualitative disease resistance and minor-gene quantitative resistance are also the common challenges of implementing and integrating any major gene or QTL into new cultivars. These difficultiesinclude inconsistent effects of the QTL due to inconsistent QTL segregations in mapping populations, QTL interaction with genetic background, and QTL interaction with the environment (Bernardo, 2008). However, in addition to the common challenges, qualitative resistance also faces the disadvantageof new virulent races of a pathogen that can over come major gene resistance (Chen and Kang, 2017). Breeding for minor gene quantitative resistance tends to produce more durable resistance in breeding lines because it relies on multiple small-effect alleles. Breeding for quantitative resistance requires multiple breeding cycles to gradually improve resistance, similar to other agronomic traits (Poland and Rutkoski, 2016). The lack of qualitative resistance durability coupled with the challenge in identifying and breeding for quantitative resistance creates a unique opportunity for genomic selection to identifyquantitative resistance by accounting for minor effect genes in the presence of large effect major genes

The goal of this study was to identify the best genomic selection method for disease resistance in the presence of both major and minor genes. In our study we used stripe rust as an example of a disease with both major and minor resistant genes. Previous studies for GS of stripe rust showed promising prediction accuracies. Muleta et al.(2017), showed that accuracy increased with population size and marker density and reached up to 0.80. Ornella et al, (2012), reported accuracies in CIMMYT wheat populations of values greater than 0.50 for stripe rust, but showed lower accuracy when compared to stem rust. Our study’s prediction accuracy for both IT and SEV reached an accuracy of up to 0.67 and 0.69 in cross-validations, respectively. Further, IT and SEV reached accuracies of up to 0.66 and 0.72 in validation sets, respectively. In comparison to other rust diseases, Rutkoski et al. (2014) and Rutkoski et al. (2015), showed promising results to predict stem rust with accuracies of up to 0.50. Overall, our study showed high prediction accuracies in comparison to most rust prediction studies, and further displayed the feasibility for accurately predicting disease resistance in the presence of major and minor resistant genes.

### Major Markers

When major genes are present, a large portion of the genetic variance for a trait may be due to unknown QTL with minor effects (Bernardo, 2014). The other minor effect QTL will not necessarily be integrated when major genes are integrated into cultivars. The lack of integration can be attributed to not being able to use MAS and the difficulties outlined previously in pyramiding major effect genes. In contrast, GS simultaneously models all QTL (Meuwissen et al., 2001). However, using GS models such as rrBLUP will underestimate the effect of the major QTL. Therefore, including the major effect QTL as fixed effects can increase accuracy. According to Bernardo (2014), major genes should be used in prediction models when only a few major genes are present, and each gene accounts for more than 10% of the variation.

In this study, the major gene in both populations was *Yr17*. In the BL, *Yr17* accounted for up to 0.40 prediction accuracy when used in MAS, and therefore accounts for a large amount of variation. The moderate accuracy of *Yr17* supports that even with the degradation of the ASR for *Yr17*, it still provides resistance for APR indicated in Liu et al. (2018). The other major rust genes present in the BL would be considered minor effect genes with near-zero prediction accuracy within MAS or, in the case of the marker for *Yr10*, only produced an accuracy above 0.10 in a few prediction scenarios. However, within the DP, all of the markers with the exception of *Lr68* produced accuracies above 0.20, with *IWB12603* reaching 0.34 and *Yr10* reaching the highest accuracies for MAS within cross-validations of 0.42, and could be considered major effect markers.

Even with the moderate accuracies of the major rust markers in MAS, we observed only a slight increase in prediction accuracy when the major markers were included in our GS models. The major markers only increased the prediction accuracy at a maximum of 0.06 within the cross-validation scenarios and 0.03 within the validation sets. Interestingly, the validation sets resulted in the highest accuracy of all scenarios with 0.72 for the base GS model and inclusion of the major markers when predicting SEV in 2014 using 2013. These results directly contrast previous studies showing higher accuracy in cross-validations (Lozada and Carter, 2019; Merrick and Carter, 2021). Validation sets are a more realistic approach for GS because it is comparable to how GS would be implemented in breeding programs (Lozada and Carter, 2019). However, the major markers only increased prediction accuracy as the overall prediction accuracy decreased. For example, using 2013-2018 to predict 2018, all of the major markers increased prediction accuracy, but the base prediction was only 0.27, and the markers increased the accuracy by 0.03 maximum in all scenarios. Further, the major markers had much larger increases in the MAS scenarios with a maximum increase of 0.17. Therefore, the inclusion of major markers provides an advantage in the more realistic validation sets when the base GS model has poor predictive ability.

In the context of GS models and breeding programs, the small increase in prediction accuracy would be considered negligible in realistic breeding scenarios. The results in our study are in contrast to previous studies that showed major markers had a large increase in prediction accuracies in GS models for other diseases such as stem rust (Rutkoski et al., 2014) and Fusarium head blight (Arruda et al., 2016). One hypothesis for the lack of increase in prediction accuracy may be due to the GS models accounting for the majority of variation in both the major and minor effect markers for disease resistance. Additionally, the lack of increase in prediction accuracy may be due to the major markers not accounting for enough phenotypic variation. The reduction in the major markers’ effect is the reason Bernardo (2014) suggested implementing markers that account for more than 10% of the variation, as mentioned previously. This theory may be disproved by the major markers displaying moderate accuracy in the MAS models. However, this may be the case for *Lr68*, which displayed minimal effect in both MAS and GS models.

Further, the lack of increase in prediction accuracy may be beneficial in demonstrating that other uncharacterized resistance QTL can still provide a large amount of disease resistance within the populations either alone or in conjunction with major genes. In this case, our results would be beneficial in confirming the presence of minor effect QTL for quantitative resistance and provide a more durable resistance within the training populations. Therefore, we can conclude that genotyping and selecting major genes for disease resistance may not be necessary when breeding programs can use more cost-effective genome-wide markers to implement GS with more consistent results.

### De Novo GWAS Markers

Frequently, the major markers for disease resistance are either unknown or have an uncharacterized effect within populations. Therefore, GWAS can be performed to characterize disease-resistant QTL within a population, and the significant markers can be used as fixed-effect covariates (Rice and Lipka, 2019). In Zhang et al. (2014), publicly available GWAS markers were integrated into prediction models but only increased the accuracy by 0.01, similar to our results. In contrast, we used de novo GWAS markers dependent on the training population. This approach has been used for FHB in which Arruda et al. (2016) demonstrated an increase in accuracy of up to 0.14. These results were also demonstrated in Spindel et al. (2016), in which de novo GWAS markers implemented into GS increased accuracies more than 0.10 in rice (*Oryza sativa* L.). However, in our study, the de novo GWAS markers only marginally increased accuracy, or in the case of implementing more than 25 markers, decreased accuracy in the majority of cross-validation scenarios. The reduction in prediction accuracy with the larger set de novo GWAS marker may be attributed to an increase in the bias of the model due to overfitting as seen in Raymond et al. (2018) or due to the difficulty for the model to simultaneously estimate all of the fixed effects (Bernardo, 2014). The reduction in prediction accuracy was also shown in Rice and Lipka (2019).

Another hypothesis for why the de novo GWAS markers failed to increase prediction accuracy may be due to the inclusion of false positives within the GWAS models. To mitigate this, we included a GWAS-GS model that only included significant markers based on a Bonferroni correction of 0.05. However, this model failed to differentiate itself from the other smaller set GWAS-GS models. The lack of reduction was mainly seen in our cross-validation sets. Within cross-validation, the training population is divided. The division of the training populations may be one cause of the lack of increase of the prediction accuracy. The smaller validation fold within cross-validation may have a weak association to the markers found in the larger training folds, as hinted at by Rice and Lipka (2019). The weak association theory may be supported by the contrasting results seen in the validation sets.

Similar to the inclusion of major markers in the cross-validations, the validation sets showed an increase in prediction accuracy when the de novo GWAS markers were included and displayed the largest increases from the GS models. The GWAS model with significant markers only (GWAS_B) displayed the largest increase of 0.06 in the SEV. Once again, this increased prediction accuracy was seen as the base GS model’s prediction accuracy decreased. This occurrence in both the major markers and de novo GWAS markers demonstrates the ability to increase and maintain high accuracy as the GS model fails in predicting lines. Therefore, we can conclude that even though fixed effect markers may not increase accuracy in typical cross-validation scenarios, they are beneficial in more realistic validation set approaches similar to major markers.

However, similar to the major markers, the increased prediction accuracy by including de novo GWAS markers was very small relative to the high accuracy for most scenarios. Further, the small sets of de novo GWAS markers were similar in consistency to the major markers. Therefore, there is little benefit in characterizing major effect disease resistance markers for GS over implementing GWAS-GS methods that would use the same sets of markers as the GS models.

### Training Population and Environment

We compared the effect of the major markers and de novo GWAS markers in different training populations that are commonly used in breeding programs. The frequency and source of both major and minor disease resistance genes vary. For instance, the BL population consists of WSU breeding lines that have been selected for resistance, specifically for *P. striiformis* f. sp. *tritici* races in Washington, and therefore, has a high level of resistance throughout the population. In comparison, the DP consists of varieties from various breeding programs in the PNW. The sources of resistance in the varieties are more similar within the BL than in the DP, with the DP containing major genes different from the major markers chosen in this study common in the WSU germplasm or selected for resistance to races not present in eastern Washington.

The differences in frequency of major genes were observed seen in the major rust markers used in this study. In the BL, the *Yr17* marker showed an increase in prediction accuracy for GS models and relatively high accuracy in the MAS models compared to the other markers. However, this was not consistently seen in the DP. The inconsistent effect of *Yr17* in different training populations may be due to the higher frequency of *Yr17* in the BL compared to the DP. This may also be supported by the higher accuracies for *Yr10* and *IWB12603* in the DP compared to the BL and both of these rust genes have a higher frequency in the DP than in the BL. Our study showed that regardless of the frequency of the rust resistant genotypes, there was only a small to zero increase in prediction accuracy. Therefore, GS would be more accurate than MAS regardless of the frequency of known rust resistant genotypes in a breeding program due to the ability to account for both major and minor disease resistance genes.

In addition to different frequencies of the major genes, the general composition of the training populations can affect GS prediction accuracy (Asoro et al., 2011). The composition of the training population affects accuracy due to both population structure and genetic relatedness (Habier et al., 2007; Asoro et al., 2011; Mirdita et al., 2015). We compared the population structure in our models by plotting principal components and identified three clusters indicating distinct subpopulations. In addition, we can see the effect of genetic relatedness and population in both our cross-validation and validation sets. The BL had a statistically higher mean accuracy for both IT and SEV than the DP in cross-validations which can be attributed to the closer genetic relatedness of the population and sources of resistance as mentioned previously. The higher prediction accuracy for the BL is advantageous for breeding programs because they can use their existing breeding trials for GS without screening a diversity panel outside of their breeding program. In the validation sets, we see an initial increase in accuracy due to the DP being the only population in the training populations, but as we added in BL lines, the accuracy decreased. The accuracy was reduced when the DP predicted the BL, but eventually increased as more BL lines were introduced into the training population. The decrease in validation sets can also be attributed to GEI (Michel et al., 2016; Huang et al., 2018; Lozada and Carter, 2019, 2020; Haile et al., 2020).

Further, GEI is important for qualitative disease resistance. Race-specific qualitative resistance is dependent on the race in the environment and thus can lead to larger environmental effects (Poland and Rutkoski, 2016). In contrast, GEI has a much smaller effect on minor-gene quantitative resistance due to the lack of gene-for-gene interaction. In our study, the most frequent races were similar from year to year, and therefore, may not be a significant factor in the differing prediction accuracy.

In this study, disease resistance screening was dependent on the natural occurrence of stripe rust for disease pressure, and therefore, the environment’s overall effect is important. Additionally, diseases such as stripe rust are affected by many environmental factors, including moisture, temperature, and wind. Further, disease severity is affected by the other aspects of the disease triangle, disease inoculum, and a susceptible host to induce disease development (Chen, 2005). Disease development may also explain the differences in prediction accuracy from year to year, especially in the DP in which the same lines are phenotyped every year. In addition, we see an increase in prediction accuracy for both the cross-validation and validation sets as we increase the number of environments within our training population. The increase in accuracy may be accounted for by the inclusion of GEI within our phenotypic adjustments and GS models as reported in previous studies, as well as the general high heritability for disease resistance (Crossa et al., 2014; Jarquín et al., 2014; Haile et al., 2020; Merrick and Carter, 2021). Overall, our GS models accurately predicted disease resistance in different training populations and environments and therefore, will be an important strategy for selecting for disease resistance.

### Applications in Breeding

Genome selection is beneficial for complex traits and can outperform phenotypic selection and MAS for low heritable traits. However, there may be little benefit in using GS for selection purposes for highly heritable traits such as disease resistance (Poland and Rutkoski, 2016). In the case of highly heritable traits, GS can still outperform phenotypic selection and MAS in terms of gain per unit time when implemented in the early stages of the breeding cycle (Bernardo and Yu, 2007; Rutkoski et al., 2011). Our study’s high prediction accuracy would allow an increase in genetic gain by decreasing the cycle time of the breeding program and rapidly accumulating favorable alleles for disease resistance (Rutkoski et al., 2011).

Even though phenotypic selection has been successfully implemented for disease resistance, without controlled experiments, one cannot determine whether the resistance is quantitative or qualitative. Therefore, we cannot conclude whether the resistance will be durable in the long term. Alternatively, we can implement MAS to select qualitative and quantitative disease resistance within the breeding lines to bypass the need for controlled experiments. However, as seen in our study, MAS does not account for all of the resistance within the lines in either of the training populations, as shown by the decrease in prediction accuracy for the MAS models. MAS also has limitations when it comes to pyramiding multiple markers, as discussed previously, and is a form of tandem selection (Bernardo, 2014). In contrast, GS is a form of selection index and has been shown to be superior to tandem selection (Hazel and Lush, 1942). Using GS, we can select for the accumulation of all resistant QTL to take advantage of the quantitative and qualitative resistant genes within a population, even when they are uncharacterized. Furthermore, by using fixed effects, we can select lines that have a major marker of interest (Poland and Rutkoski, 2016). Therefore, GS will have a place in selecting for both quantitative and qualitative disease resistance.

Another advantage in implementing GS is by reducing both genotyping and phenotyping within a breeding program. GS can remove the need for genotyping for major and minor genes for selection purposes. This is further supported by the similar accuracies between major and de novo GWAS markers. By utilizing genome-wide markers, we can not only implement GS or GWAS-GS but also utilize the markers for additional traits, thus making the genome-wide markers more cost-effective (Poland and Rutkoski, 2016). Likewise, by using GS, breeding programs can reduce the need for phenotypic screening in disease nurseries in multiple locations and free up resources for screening more lines and increase genetic gain (Poland and Rutkoski, 2016).

Furthermore, the challenges introduced by the environment mentioned previously provide another advantage in using GS for disease resistance. The GS models will help select cultivars with durable quantitative resistance with the accumulation of favorable alleles and select for disease resistance in environments not conducive to disease incidence needed for phenotypic selection. Overall, the high accuracy of the GS models in our study displays the ability to predict durable disease resistance and account for uncharacterized minor effect QTL in the presence of known major genes.

## CONCLUSIONS

This study showed the ability to predict disease resistance using major and minor genes accurately. The small to no increase in prediction accuracy using major markers indicates the need for careful selection of major markers that account for a large variation in the training and test populations. Further, the comparison of the number of de novo GWAS markers shows a small number of de novo GWAS markers should be used instead of a large set of markers to keep from overfitting the model. Additionally, fixed effect markers may not provide a benefit in scenarios with already high prediction accuracy. However, in prediction scenarios with low accuracies, such as in more realistic validation sets, the inclusion of both major markers and de novo GWAS helps account for variation when the base GS models fail. Moreover, we can increase accuracy with the inclusion of additional environments and by using populations that are genetically related. Overall, there were no disadvantages in the inclusion of the major markers or de novo GWAS markers. The lack of increase of prediction accuracy with the inclusion of fixed effects coupled with the large decrease in accuracy using MAS indicates the presence of minor effect QTL for quantitative resistance and thus durable resistance within the training populations. This study showed the ability to predict disease resistance and accumulate favorable alleles for durable disease resistance in the presence of major and minor resistance genes.

## DATA AVAILABILITY STATEMENT

The datasets generated for this study can be found at https://github.com/lfmerrick21/Major-and-Minor-Genes

## AUTHOR CONTRIBUTIONS

LM: conceptualized the idea, analysed data, and drafted the manuscript; AB: genotyped the KASP markers, reviewed, and edited the manuscript; XC: reviewed and edited the manuscript; AC: supervised the study, conducted field trials, edited the manuscript and obtained the funding for the project.

## ACKNOWLEDGMENTS

The authors would like to acknowledge the Washington State University Winter Wheat Breeding Program personnel Gary Shelton and Kyall Hagemeyer for plot maintenance and data collection under field conditions. We would also like to thank Gina Brown-Guedira, Jared Smith, Brian Ward and staff at the Eastern Regional Small Grains Genotyping Laboratory for their assistance with DNA library prep and GBS sequencing and analysis.

## SUPPLEMENTARY MATERIAL

Supplementary tables are available in a separate file.

