## Supplementary Material for "Breeding with Major and Minor Genes: Genomic Selection for Quantitative Disease Resistance"

**Table S1.** Stripe rust race summary and major race frequency across Eastern Washington in 2013 to 2020.

| Year | No. of Isolates | No. of races | Major Races | Freqency (%) |
|---|---|---|---|---|
| 2020 | 158 | 14 | PSTv-37 | 50.6 |
| | | | PSTv-39 | 22.8 |
| 2018 | 150 | 20 | PSTv-37 | 38.7 |
| | | | PSTv-52 | 10.7 |
| | | | PSTv-322 | 8 |
| 2017 | 104 | 43 | PSTv-37 | 40.4 |
| | | | PSTv-322 | 4.8 |
| 2016 | 149 | 37 | PSTv-37 | 19.5 |
| | | | PSTv-52 | 13.4 |
| 2015 | 130 | 23 | PSTv-37 | 30 |
| | | | PSTv-52 | 23.8 |
| 2014 | 129 | 22 | PSTv-37 | 10.1 |
| | | | PSTv-48 | 10.9 |
| | | | PSTv-52 | 25.6 |
| | | | PSTv-79 | 14.7 |
| 2013 | 180 | 26 | PSTv-11 | 10 |
| | | | PSTv-37 | 12.8 |
| | | | PSTv-52 | 26.7 |
| | | | PSTv-73 | 9.4 |

**Table S2.** Comparison of genomic selection models accuracy and pairwise comparisons for stripe rust infection type for Pacific Northwest winter wheat diversity panel (DP) lines and breeding lines (BL) phenotyped from 2013-2020 in Central Ferry, Lind, and Pullman, WA using cross-validation within and across trials.

| Population | Year | Reg | IWB12603 | *Lr68* | *Yr10* | *Yr17* | All_M | GWAS_B | GWAS_5 | GWAS_10 | GWAS_25 | GWAS_50 | GWAS_100 |
|---|---|---|---|---|---|---|---|---|---|---|---|---|---|
| BL | 2016 | 0.57cd | 0.56d | 0.57cd | 0.58bc | 0.60ab | 0.61a | 0.53e | 0.50f | 0.48g | 0.46h | 0.45h | 0.42i |
| BL | 2017 | 0.48bc | 0.48bc | 0.48bc | 0.47cd | 0.49a | 0.48ab | 0.46d | 0.46d | 0.43e | 0.40f | 0.39fg | 0.38g |
| BL | 2018 | 0.65cd | 0.65d | 0.65cd | 0.65cd | 0.65bcd | 0.66bcd | 0.66b | 0.67a | 0.66bc | 0.63e | 0.62f | 0.60g |
| BL | 2020 | 0.65bc | 0.67ab | 0.66ab | 0.66ab | 0.66ab | 0.67a | 0.66ab | 0.66ab | 0.64c | 0.61d | 0.59e | 0.56f |
| BL | 2016-2017 | 0.51b | 0.51b | 0.51b | 0.51b | 0.52a | 0.52a | 0.49c | 0.49c | 0.47d | 0.45e | 0.43f | 0.43f |
| BL | 2016-2018 | 0.60c | 0.61c | 0.60c | 0.60c | 0.61ab | 0.61ab | 0.61bc | 0.61a | 0.61a | 0.60d | 0.58e | 0.56f |
| BL | 2016-2020 | 0.61e | 0.61e | 0.61de | 0.61de | 0.62bc | 0.61bc | 0.61cd | 0.62a | 0.62ab | 0.60f | 0.59g | 0.58h |
| DP | 2013 | 0.55a | 0.55a | 0.55a | 0.55a | 0.55a | 0.55a | 0.47b | 0.48b | 0.45c | 0.43d | 0.40e | 0.41e |
| DP | 2014 | 0.46a | 0.45a | 0.45a | 0.46a | 0.45a | 0.44a | 0.40b | 0.39b | 0.35cd | 0.34cd | 0.35c | 0.34d |
| DP | 2015 | 0.54a | 0.54a | 0.54a | 0.55a | 0.54a | 0.54a | 0.49b | 0.48b | 0.45c | 0.43d | 0.41d | 0.39e |
| DP | 2016 | 0.5a | 0.49a | 0.50a | 0.50a | 0.49a | 0.49a | 0.41b | 0.41b | 0.36c | 0.35cd | 0.34de | 0.33e |
| DP | 2013-2014 | 0.54a | 0.54a | 0.54a | 0.55a | 0.54a | 0.54a | 0.47b | 0.48b | 0.44c | 0.42cd | 0.41de | 0.40e |
| DP | 2013-2015 | 0.54a | 0.54a | 0.54a | 0.55a | 0.55a | 0.54a | 0.52b | 0.51b | 0.50c | 0.47d | 0.47d | 0.46d |
| DP | 2013-2016 | 0.56a | 0.56a | 0.57a | 0.57a | 0.56a | 0.56a | 0.54b | 0.53b | 0.51c | 0.49d | 0.48de | 0.47e |

Models labeled with the same letter are not significantly different (*P*-value =0.05). Adjustments: ALL_M: IWB12603(Qyr.wpg-1B.1), KASP(Lr68), Xpsp3000(Yr10), and KASP(Yr17) combined; GWAS_B: genome-wide association assisted genomic selection (GWAS-GS) with Bonferonni significant markers; GWAS_5: GWAS-GS with the top 5 significant markers; GWAS_10: GWAS-GS with the top 10 significant markers; GWAS_25: GWAS-GS with the top 25 significant markers; GWAS_50: GWAS-GS with the top 50 significant markers; and GWAS_100: GWAS-GS with the top 100 significant markers.

**Table S3.** Comparison of genomic selection models accuracy and pairwise comparisons for stripe rust disease severity for Pacific Northwest winter wheat diversity panel (DP) lines and breeding lines (BL) phenotyped from 2013-2020 in Central Ferry, Lind, and Pullman, WA using cross-validation within and across trials.

| Population | Year | Reg | IWB12603 | *Lr68* | *Yr10* | *Yr17* | All_M | GWAS_B | GWAS_5 | GWAS_10 | GWAS_25 | GWAS_50 | GWAS_100 |
|---|---|---|---|---|---|---|---|---|---|---|---|---|---|
| BL | 2016 | 0.51c | 0.53bc | 0.52c | 0.55b | 0.54bc | 0.58a | 0.40d | 0.40d | 0.38d | 0.35e | 0.34ef | 0.32f |
| BL | 2017 | 0.50bc | 0.50c | 0.50cd | 0.50cd | 0.50abc | 0.50abc | 0.51a | 0.51ab | 0.49d | 0.47e | 0.45f | 0.44g |
| BL | 2018 | 0.68bc | 0.68bc | 0.68bc | 0.68bc | 0.68bc | 0.68bc | 0.68b | 0.69a | 0.67c | 0.66d | 0.64e | 0.63f |
| BL | 2020 | 0.66ab | 0.67a | 0.66ab | 0.67a | 0.67a | 0.66ab | 0.64cd | 0.65bc | 0.63d | 0.61e | 0.60f | 0.57g |
| BL | 2016-2017 | 0.51bc | 0.51bc | 0.52abc | 0.52ab | 0.52ab | 0.53a | 0.51cd | 0.51bc | 0.50d | 0.48e | 0.46f | 0.45g |
| BL | 2016-2018 | 0.63c | 0.63c | 0.63c | 0.63c | 0.63c | 0.63c | 0.64b | 0.64ab | 0.64a | 0.63c | 0.62d | 0.60e |
| BL | 2016-2020 | 0.63bcd | 0.63cd | 0.63bcd | 0.63bcd | 0.63b | 0.63bc | 0.63a | 0.64a | 0.64a | 0.62d | 0.61e | 0.60f |
| DP | 2013 | 0.64a | 0.64a | 0.64a | 0.65a | 0.64a | 0.64a | 0.59b | 0.59b | 0.56c | 0.53d | 0.52e | 0.52e |
| DP | 2014 | 0.63a | 0.63a | 0.63a | 0.63a | 0.63a | 0.63a | 0.59b | 0.61b | 0.58c | 0.55d | 0.54de | 0.53e |
| DP | 2015 | 0.60ab | 0.59b | 0.6ab | 0.60a | 0.59ab | 0.6ab | 0.56c | 0.55d | 0.51e | 0.48f | 0.47g | 0.46g |
| DP | 2016 | 0.58a | 0.57a | 0.58a | 0.58a | 0.58a | 0.57a | 0.55b | 0.54b | 0.51c | 0.49d | 0.48de | 0.47e |
| DP | 2013-2014 | 0.69a | 0.68a | 0.68a | 0.69a | 0.68a | 0.69a | 0.65c | 0.67b | 0.63d | 0.61e | 0.60e | 0.58f |
| DP | 2013-2015 | 0.65a | 0.65a | 0.65a | 0.66a | 0.65a | 0.65a | 0.61b | 0.61b | 0.59c | 0.57d | 0.57d | 0.56e |
| DP | 2013-2016 | 0.67a | 0.67a | 0.67a | 0.67a | 0.67a | 0.67a | 0.63c | 0.64b | 0.61d | 0.58e | 0.58ef | 0.57f |

Models labeled with the same letter are not significantly different (*P*-value =0.05). Adjustments: ALL_M: IWB12603(Qyr.wpg-1B.1), KASP(Lr68), Xpsp3000(Yr10), and KASP(Yr17) combined; GWAS_B: genome-wide association assisted genomic selection (GWAS-GS) with Bonferonni significant markers; GWAS_5: GWAS-GS with the top 5 significant markers; GWAS_10: GWAS-GS with the top 10 significant markers; GWAS_25: GWAS-GS with the top 25 significant markers; GWAS_50: GWAS-GS with the top 50 significant markers; and GWAS_100: GWAS-GS with the top 100 significant markers.

**Table S4.** Comparison of marker-assisted selection models accuracy and pairwise comparisons for stripe rust infection type for Pacific Northwest winter wheat diversity panel (DP) lines and breeding lines (BL) phenotyped from 2013-2020 in Central Ferry, Lind, and Pullman, WA using cross-validation within and across trials.

| Population | Year | IWB12603 | *Lr68* | *Yr10* | *Yr17* | All_M | GWAS_B | GWAS_5 | GWAS_10 | GWAS_25 | GWAS_50 | GWAS_100 |
|---|---|---|---|---|---|---|---|---|---|---|---|---|
| BL | 2016 | 0.05c | 0.03c | -0.01d | 0.40ab | 0.40ab | 0.39ab | 0.38b | 0.41a | 0.41a | 0.41a | 0.41a |
| BL | 2017 | 0.06e | 0.15d | 0.15d | 0.21c | 0.29b | 0.36a | 0.37a | 0.38a | 0.36a | 0.37a | 0.37a |
| BL | 2018 | 0.09f | -0.03h | 0.07g | 0.45e | 0.45e | 0.60a | 0.59bc | 0.60a | 0.60ab | 0.59cd | 0.58d |
| BL | 2020 | 0.10e | 0.07f | 0.08ef | 0.25d | 0.24d | 0.45c | 0.46c | 0.50b | 0.52ab | 0.53a | 0.52a |
| BL | 2016-2017 | 0.04g | 0.08f | 0.12e | 0.27d | 0.30c | 0.36b | 0.37b | 0.39a | 0.39a | 0.39a | 0.39a |
| BL | 2016-2018 | 0.04h | 0.03i | 0.08g | 0.38f | 0.39e | 0.51b | 0.48d | 0.51c | 0.54a | 0.54a | 0.54a |
| BL | 2016-2020 | 0.01j | 0.02i | 0.08h | 0.36g | 0.37f | 0.52c | 0.46e | 0.51d | 0.54b | 0.55a | 0.55a |
| DP | 2013 | 0.31d | 0.08h | 0.38b | 0.19g | 0.42a | 0.19g | 0.23f | 0.27e | 0.32d | 0.35c | 0.35c |
| DP | 2014 | 0.14f | -0.07g | 0.23c | 0.21d | 0.28b | 0.19e | 0.20de | 0.25c | 0.28b | 0.30ab | 0.31a |
| DP | 2015 | 0.33d | 0.08g | 0.42b | 0.16f | 0.44a | 0.30e | 0.33d | 0.37c | 0.38c | 0.39c | 0.38c |
| DP | 2016 | 0.21d | 0.01g | 0.32a | 0.19e | 0.35a | 0.15f | 0.18e | 0.23d | 0.26c | 0.29b | 0.29b |
| DP | 2013-2014 | 0.23e | 0.01g | 0.34b | 0.21ef | 0.38a | 0.21ef | 0.20f | 0.26d | 0.31c | 0.36ab | 0.36ab |
| DP | 2013-2015 | 0.33f | 0.09h | 0.40d | 0.15g | 0.42c | 0.37e | 0.38de | 0.42c | 0.43bc | 0.44ab | 0.44a |
| DP | 2013-2016 | 0.34e | 0.08g | 0.40c | 0.16f | 0.43ab | 0.38d | 0.38d | 0.42bc | 0.44a | 0.44a | 0.45a |

Models labeled with the same letter are not significantly different (*P*-value =0.05). Adjustments: ALL_M: IWB12603(Qyr.wpg-1B.1), KASP(Lr68), Xpsp3000(Yr10), and KASP(Yr17) combined; GWAS_B: genome-wide association assisted genomic selection (GWAS-GS) with Bonferonni significant markers; GWAS_5: GWAS-GS with the top 5 significant markers; GWAS_10: GWAS-GS with the top 10 significant markers; GWAS_25: GWAS-GS with the top 25 significant markers; GWAS_50: GWAS-GS with the top 50 significant markers; and GWAS_100: GWAS-GS with the top 100 significant markers.

**Table S5.** Comparison of marker-assisted selection models accuracy and pairwise comparisons for stripe rust disease severity for Pacific Northwest winter wheat diversity panel (DP) lines and breeding lines (BL) phenotyped from 2013-2020 in Central Ferry, Lind, and Pullman, WA using cross-validation within and across trials.

| Population | Year | IWB12603 | *Lr68* | *Yr10* | *Yr17* | All_M | GWAS_B | GWAS_5 | GWAS_10 | GWAS_25 | GWAS_50 | GWAS_100 |
|---|---|---|---|---|---|---|---|---|---|---|---|---|
| BL | 2016 | 0.17c | -0.04d | -0.09e | 0.42a | 0.43a | 0.24b | 0.22b | 0.23b | 0.25b | 0.24b | 0.24b |
| BL | 2017 | 0.10i | 0.15h | 0.19g | 0.23f | 0.33e | 0.44ab | 0.45a | 0.44ab | 0.43bc | 0.43cd | 0.42d |
| BL | 2018 | 0.09f | 0.00h | 0.07g | 0.45e | 0.46d | 0.63a | 0.63a | 0.63a | 0.62a | 0.61b | 0.61c |
| BL | 2020 | -0.05g | -0.07g | -0.04g | 0.22e | 0.19f | 0.40d | 0.38d | 0.44c | 0.48b | 0.51ab | 0.52a |
| BL | 2016-2017 | 0.05i | 0.10h | 0.14g | 0.28f | 0.33e | 0.44bc | 0.41d | 0.45a | 0.44ab | 0.44abc | 0.43c |
| BL | 2016-2018 | 0.03i | 0.05h | 0.10g | 0.39f | 0.40e | 0.57b | 0.51d | 0.56c | 0.58a | 0.58a | 0.58a |
| BL | 2016-2020 | 0.02j | 0.03i | 0.08h | 0.37g | 0.38f | 0.55c | 0.49e | 0.53d | 0.57b | 0.57a | 0.58a |
| DP | 2013 | 0.34f | 0.06h | 0.43d | 0.29g | 0.51a | 0.35f | 0.37e | 0.42d | 0.46c | 0.47bc | 0.48b |
| DP | 2014 | 0.38f | -0.06g | 0.42e | 0.38f | 0.56a | 0.46d | 0.48cd | 0.49c | 0.51b | 0.51b | 0.52b |
| DP | 2015 | 0.33e | -0.04g | 0.43b | 0.27f | 0.49a | 0.34e | 0.38d | 0.40c | 0.43b | 0.43b | 0.44b |
| DP | 2016 | 0.33f | -0.08g | 0.39d | 0.34e | 0.51a | 0.41d | 0.43c | 0.45b | 0.47b | 0.45b | 0.45b |
| DP | 2013-2014 | 0.38e | -0.07g | 0.46d | 0.36f | 0.57a | 0.49c | 0.48c | 0.53b | 0.56a | 0.56a | 0.57a |
| DP | 2013-2015 | 0.41d | 0.03f | 0.46c | 0.28e | 0.54ab | 0.47c | 0.47c | 0.52b | 0.54a | 0.55a | 0.54a |
| DP | 2013-2016 | 0.42e | 0.01g | 0.46d | 0.30f | 0.56a | 0.49c | 0.50c | 0.53b | 0.55a | 0.56a | 0.55a |

Models labeled with the same letter are not significantly different (*P*-value =0.05). Adjustments: ALL_M: IWB12603 IWB12603(Qyr.wpg-1B.1), KASP(Lr68), Xpsp3000(Yr10), and KASP(Yr17) combined; GWAS_B: genome-wide association assisted genomic selection (GWAS-GS) with Bonferonni significant markers; GWAS_5: GWAS-GS with the top 5 significant markers; GWAS_10: GWAS-GS with the top 10 significant markers; GWAS_25: GWAS-GS with the top 25 significant markers; GWAS_50: GWAS-GS with the top 50 significant markers; GWAS_100: GWAS-GS with the top 100 significant markers.

**Table S6.** Comparison of genomic selection models accuracy and pairwise comparisons for stripe rust infection type for Pacific Northwest winter wheat diversity panel (DP) lines and breeding lines (BL) phenotyped from 2013-2020 in Central Ferry, Lind, and Pullman, WA using validation sets.

| Validation Set | Reg | IWB12603 | *Lr68* | *Yr10* | *Yr17* | All_M | GWAS_B | GWAS_5 | GWAS_10 | GWAS_25 | GWAS_50 | GWAS_100 |
|---|---|---|---|---|---|---|---|---|---|---|---|---|
| 2013 to 2014 | 0.56d | 0.57c | 0.57b | 0.56f | 0.57a | 0.56e | 0.55g | 0.54j | 0.54k | 0.53l | 0.54h | 0.54i |
| 2013-2014 to 2015 | 0.65d | 0.65e | 0.65c | 0.66b | 0.65f | 0.66a | 0.65i | 0.65h | 0.65g | 0.62j | 0.62k | 0.61l |
| 2013-2015 to 2016 | 0.53c | 0.52f | 0.53a | 0.53d | 0.53b | 0.52e | 0.52g | 0.50h | 0.49i | 0.42l | 0.44k | 0.45j |
| 2013-2016 to 2017 | 0.37f | 0.37e | 0.36h | 0.37g | 0.38d | 0.38c | 0.39b | 0.39a | 0.35i | 0.29j | 0.25k | 0.25l |
| 2013-2017 to 2018 | 0.27j | 0.28i | 0.28h | 0.28g | 0.31e | 0.31f | 0.32b | 0.33a | 0.32c | 0.31d | 0.27k | 0.23l |
| 2013-2018 to 2020 | 0.51k | 0.52g | 0.51j | 0.52h | 0.52f | 0.52e | 0.55a | 0.53d | 0.54b | 0.54c | 0.50l | 0.51i |

Models labeled with the same letter are not significantly different (*P*-value =0.05). Adjustments: ALL_M: IWB12603(Qyr.wpg-1B.1), KASP(Lr68), Xpsp3000(Yr10), and KASP(Yr17) combined; GWAS_B: genome-wide association assisted genomic selection (GWAS-GS) with Bonferonni significant markers; GWAS_5: GWAS-GS with the top 5 significant markers; GWAS_10: GWAS-GS with the top 10 significant markers; GWAS_25: GWAS-GS with the top 25 significant markers; GWAS_50: GWAS-GS with the top 50 significant markers; and GWAS_100: GWAS-GS with the top 100 significant markers.

**Table S7.** Comparison of genomic selection models accuracy and pairwise comparisons for stripe rust disease severity for Pacific Northwest winter wheat diversity panel (DP) lines and breeding lines (BL) phenotyped from 2013-2020 in Central Ferry, Lind, and Pullman, WA using validation sets.

| Validation Set | Reg | IWB12603 | *Lr68* | *Yr10* | *Yr17* | All_M | GWAS_B | GWAS_5 | GWAS_10 | GWAS_25 | GWAS_50 | GWAS_100 |
|---|---|---|---|---|---|---|---|---|---|---|---|---|
| 2013 to 2014 | 0.72e | 0.72d | 0.72c | 0.72f | 0.72b | 0.72a | 0.71h | 0.71g | 0.70i | 0.69l | 0.69k | 0.69j |
| 2013-2014 to 2015 | 0.70h | 0.70g | 0.70f | 0.71d | 0.7e | 0.71a | 0.71b | 0.71c | 0.70i | 0.69j | 0.69k | 0.68l |
| 2013-2015 to 2016 | 0.64b | 0.63d | 0.64c | 0.63e | 0.64a | 0.63f | 0.59g | 0.58h | 0.57i | 0.52l | 0.54k | 0.55j |
| 2013-2016 to 2017 | 0.42d | 0.42e | 0.41g | 0.42b | 0.42a | 0.42c | 0.38j | 0.38k | 0.40h | 0.41f | 0.38i | 0.37l |
| 2013-2017 to 2018 | 0.32l | 0.33j | 0.33g | 0.33i | 0.34e | 0.34f | 0.38a | 0.33h | 0.37b | 0.35c | 0.35d | 0.32k |
| 2013-2018 to 2020 | 0.47j | 0.47h | 0.47i | 0.46k | 0.44l | 0.51a | 0.5b | 0.47f | 0.50c | 0.47e | 0.47g | 0.48d |

Models labeled with the same letter are not significantly different (*P*-value =0.05). Adjustments: ALL_M: IWB12603(Qyr.wpg-1B.1), KASP(Lr68), Xpsp3000(Yr10), and KASP(Yr17) combined; GWAS_B: genome-wide association assisted genomic selection (GWAS-GS) with Bonferonni significant markers; GWAS_5: GWAS-GS with the top 5 significant markers; GWAS_10: GWAS-GS with the top 10 significant markers; GWAS_25: GWAS-GS with the top 25 significant markers; GWAS_50: GWAS-GS with the top 50 significant markers; and GWAS_100: GWAS-GS with the top 100 significant markers.

**Table S8.** Comparison of marker-assisted selection models accuracy and pairwise comparisons for stripe rust infection type for Pacific Northwest winter wheat diversity panel (DP) lines and breeding lines (BL) phenotyped from 2013-2020 in Central Ferry, Lind, and Pullman, WA using validation sets.

| Validation Set | IWB12603 | *Lr68* | *Yr10* | *Yr17* | All_M | GWAS_B | GWAS_5 | GWAS_10 | GWAS_25 | GWAS_50 | GWAS_100 |
|---|---|---|---|---|---|---|---|---|---|---|---|
| 2013 to 2014 | 0.14j | 0.01k | 0.24h | 0.21i | 0.29g | 0.32f | 0.33e | 0.41d | 0.47c | 0.50b | 0.56a |
| 2013-2014 to 2015 | 0.33f | 0.08k | 0.42e | 0.17i | 0.44d | 0.13j | 0.24h | 0.26g | 0.56c | 0.57b | 0.62a |
| 2013-2015 to 2016 | 0.05i | -0.03k | 0.12h | 0.33c | 0.23e | 0.20f | 0.20g | 0.30d | 0.35b | 0.37a | 0.04j |
| 2013-2016 to 2017 | 0.06i | -0.15j | 0.15g | 0.21f | 0.24e | 0.38a | 0.38a | 0.30b | 0.26c | 0.24d | 0.12h |
| 2013-2017 to 2018 | -0.09k | -0.01j | 0.07i | 0.45b | 0.43d | 0.43c | 0.47a | 0.39e | 0.30f | 0.25g | 0.23h |
| 2013-2018 to 2020 | -0.10k | 0.07j | 0.08i | 0.25h | 0.27g | 0.51a | 0.47d | 0.46e | 0.50b | 0.50c | 0.42f |

Models labeled with the same letter are not significantly different (*P*-value =0.05). Adjustments: ALL_M: IWB12603(Qyr.wpg-1B.1), KASP(Lr68), Xpsp3000(Yr10), and KASP(Yr17) combined; GWAS_B: genome-wide association assisted genomic selection (GWAS-GS) with Bonferonni significant markers; GWAS_5: GWAS-GS with the top 5 significant markers; GWAS_10: GWAS-GS with the top 10 significant markers; GWAS_25: GWAS-GS with the top 25 significant markers; GWAS_50: GWAS-GS with the top 50 significant markers; and GWAS_100: GWAS-GS with the top 100 significant markers.

**Table S9.** Comparison of marker-assisted selection models accuracy and pairwise comparisons for stripe rust disease severity for Pacific Northwest winter wheat diversity panel (DP) lines and breeding lines (BL) phenotyped from 2013-2020 in Central Ferry, Lind, and Pullman, WA using validation sets.

| Validation Set | IWB12603 | *Lr68* | *Yr10* | *Yr17* | All_M | GWAS_B | GWAS_5 | GWAS_10 | GWAS_25 | GWAS_50 | GWAS_100 |
|---|---|---|---|---|---|---|---|---|---|---|---|
| 2013 to 2014 | 0.37j | -0.01k | 0.42h | 0.38i | 0.56f | 0.56e | 0.55g | 0.60d | 0.63c | 0.65b | 0.68a |
| 2013-2014 to 2015 | 0.33i | 0.02k | 0.43h | 0.27j | 0.50g | 0.55e | 0.51f | 0.59d | 0.59c | 0.64b | 0.68a |
| 2013-2015 to 2016 | 0.12i | -0.05k | 0.15h | 0.41e | 0.38f | 0.48b | 0.26g | 0.47c | 0.46d | 0.50a | 0.07j |
| 2013-2016 to 2017 | 0.10i | -0.15j | 0.19f | 0.23e | 0.29d | 0.11h | 0.11h | 0.30c | 0.34b | 0.34a | 0.13g |

| | | | | | | | | | | | |
|---|---|---|---|---|---|---|---|---|---|---|---|
| 2013-2017 to 2018 | -0.09k | 0.02j | 0.07i | 0.45a | 0.42c | 0.43b | 0.08h | 0.36e | 0.36d | 0.35f | 0.18g |
| 2013-2018 to 2020 | 0.02j | 0.02k | 0.02i | 0.23g | 0.22h | 0.41e | 0.41d | 0.47a | 0.41f | 0.45b | 0.43c |

Models labeled with the same letter are not significantly different (*P*-value =0.05). Adjustments: ALL_M: IWB12603(Qyr.wpg-1B.1), KASP(Lr68), Xpsp3000(Yr10), and KASP(Yr17) combined; GWAS_B: genome-wide association assisted genomic selection (GWAS-GS) with Bonferonni significant markers; GWAS_5: GWAS-GS with the top 5 significant markers; GWAS_10: GWAS-GS with the top 10 significant markers; GWAS_25: GWAS-GS with the top 25 significant markers; GWAS_50: GWAS-GS with the top 50 significant markers; and GWAS_100: GWAS-GS with the top 100 significant markers.
